# Genome-scale multi-organ analysis of mutagenic effects of ethanol and acetaldehyde in Sprague Dawley rats

**DOI:** 10.1101/2025.09.01.673482

**Authors:** Bérénice Chavanel, Eva Tibaldi, François Virard, Christine Carreira, Sergey Senkin, Behnoush Abedi-Ardekani, Vincent Cahais, Michael Korenjak, Zdenko Herceg, Silvia Balbo, Daniele Mandrioli, Jiri Zavadil

## Abstract

Alcohol consumption is a major cancer risk factor, particularly for head and neck cancers, including the oral cavity. Acetaldehyde, the primary genotoxic metabolite of ethanol, may play key roles in oral carcinogenesis, though the mechanisms remain unclear. While mutational signatures SBS16, DBS4, and ID11 have been tentatively linked to alcohol use, they are not exclusive to alcohol-related cancers.

In this study, we examined the genome-wide *in vivo* mutagenic effects of ethanol and acetaldehyde by analyzing tumors from the cheek, Zymbal gland, larynx, forestomach, and liver of rats chronically exposed to these compounds. Signature analysis revealed exposure-specific, early-onset formation of SBS17 in ∼28% of head and neck tumors, suggesting inflammation and/or oxidative damage as potential mediators of carcinogenesis. Cancer driver gene analysis identified a relative enrichment of exposed tumors with mutations in the *Tp53* and *Mtor* genes. No notable exposure-specific changes were observed in doublet-base substitutions, indel signatures, or copy number variants. Notably, SBS16, DBS4, and ID11 were absent. Our findings suggest direct mutagenicity may not be the main driver of alcohol-related cancer. Other harmful cellular effects, undetectable by whole genome sequencing, may be involved. Our findings suggest that SBS17 could function as a potential exposure-specific molecular marker of alcohol-related cancers in humans.

## INTRODUCTION

Alcohol drinking is responsible for numerous diseases and health conditions and is one of the major risk factors for developing cancer at various anatomical sites. Ethanol (EtOH) and its primary metabolite, acetaldehyde (AcA), in alcoholic beverages are both classified by the IARC Monographs program as carcinogenic to humans (i.e. Group 1 carcinogen) [1]. Alcohol intake increases the risk of at least eight different cancer types, including the oral cavity, pharynx, larynx, esophagus, liver, breast, pancreas and colorectum, and together, these cancers contributed ∼4.1% of all new cancer cases worldwide in 2020 [2]. The mechanisms implicated in the carcinogenicity of alcohol are diverse and not entirely characterized, although AcA may play a primary role in alcohol-related carcinogenesis. EtOH and its byproduct AcA exhibit well-documented genotoxic and mutagenic effects that are believed to significantly contribute to their role in carcinogenesis and cellular dysfunction, inducing various types of DNA damage, including single- and double-strand DNA breaks, bulky adducts, and inter-and intra-strand crosslinks, as well as point mutations [3–5].

Deciphering mutagenesis processes caused by a certain chemical exposure or specific endogenous pathway can be approached through the exploration of the induced mutational signatures [6–12]. As numerous of these signatures are associated with specific mutagenic processes, their presence provides evidence that a given process plays a role in carcinogenesis [13–17]. The sources of mutational signatures are either endogenous and occurring naturally in the cells (e.g. DNA repair deficiency, deregulation of intrinsic mutagenic enzymes), or result from exogenous exposures to mutagens coming from environmental, lifestyle or therapeutic sources, such as tobacco smoking, asbestos exposure or UV light. The exploration and understanding of the causes of mutational signatures are particularly useful in cancer research as they provide clues about the biological mechanisms, environmental factors, or exposures that contributed to the initiation and development of cancer.

Although alcohol is a well-established human carcinogen, no mutational signature has been unambiguously associated with its consumption thus far. COSMIC signature SBS16 has been tentatively linked to alcohol drinking and is observed most frequently in liver cancer cases [18]. Accumulation of SBS16-specific mutations was found in alcohol-drinking patients with liver tumors [19, 20] and esophageal squamous cell carcinomas [21–23]. A correlation between the inactivity of the AcA-detoxifying enzyme ALDH2 and the formation of SBS16 was observed in ALDH2 deficient individuals, suggesting a role for AcA derived from ethanol metabolism in inducing the SBS16-specific mutation pattern [21, 22, 24]. However, SBS16 has also been found in the absence of alcohol exposure history [25–28]. Other studies observed the formation of the signature in the context of exposure to tobacco smoking [29]. SBS16 mutations were specifically identified in current and ex-smoker samples with a dose-response induction of the signature with increasing pack-years of tobacco use [30, 31]. More recently, the double-base substitution signature DBS4 and the indel signature ID11 were found, in addition to SBS16, in genomes of head and neck [32, 33] and liver [34] cancers of drinkers, but their presence was strongly enriched upon co-exposure to alcohol and tobacco smoking [32, 33, 35–37]. This raises the question of whether alcohol itself promotes specific mutagenesis or rather acts in combination with other carcinogens such as tobacco, by enhancing the effects of the latter. In addition, some studies proposed that alcohol exposure has no direct mutagenic effect, and thus is not associated with the formation of SBS16 or other mutational signatures. Rather, chronic alcohol consumption was proposed to affect liver stem cells prior to cancer development, resulting in microenvironment changes that are permissive for the selection of cells with oncogenic mutations, such as the induction of a chronic inflammation field or an immunosuppressive microenvironment that potentially drives the transition from healthy to precancerous stem cells and enhances liver tumorigenesis [38, 39]. Previous attempts using *in vitro* experimental systems to investigate the mutagenic effects of EtOH or AcA and to link these exposures to a defined mutational process or signature remain inconclusive in generating a reproducible AcA-derived mutational signature in iPS cells [9] or in yeast models [40, 41], emphasizing that solid experimental evidence for specific effects of EtOH and AcA on genome-wide mutagenesis remains lacking.

Studies of the effects of chemical compounds in rodents provide invaluable insights into how these agents initiate and promote cancer development, offering an approximation of the human physiological context. Rodent models are commonly used in chemical carcinogenicity bioassays to evaluate potential human carcinogens [42–44], due to the similarity of relevant biological pathways between rodents and humans. An additional advantage of rodent bioassays is their ability to capture the metabolization of chemicals *in vivo*, a feature that often lacks in cell-based systems, which may not fully replicate the complex interactions of metabolic activation and detoxification pathways. Studies showed that rodent models reproduce mutagenic patterns induced in other experimental settings and found in human cancer data [45, 46], and also enabled assessing the effects of non-genotoxic carcinogens acting as cancer promoters [47]. Importantly, the carcinogen evaluation and classification program conducted by the IARC Monographs considers the outcomes from animal bioassays as a key evidence stream.

Experimental studies conducted in mice by the U.S. National Toxicology Program assessed the toxicological and carcinogenic effects of ethanol in animals, with inconclusive findings regarding a link between exposure through drinking water and cancer development [48]. In contrast, Sprague Dawley rat exposure bioassays performed at the Ramazzini Institute, Bologna, Italy, as part of the long-term experimental studies on the carcinogenicity of EtOH [49] and AcA [50] indicated organ-specific tumorigenic effects for either exposure. We thus hypothesized that direct DNA damage by EtOH/AcA can be effectively assessed in archived tissue samples derived from these studies performed in well-controlled experimental settings. In this study, we analyzed the DNA mutational patterns in tumors formed in Sprague Dawley rats chronically exposed to EtOH or AcA through drinking water. By performing whole genome sequencing of the rat tumors, and through the analysis of mutational signatures, copy number alterations and EtOH/AcA-associated mutation burdens in cancer driver genes, we aimed to comprehensively describe the *in vivo* mutational underpinnings of alcohol-related carcinogenesis.

## MATERIALS AND METHODS

### Rat tissues

This study analyzed alcohol-fixed paraffin-embedded (AFPE) tumor and non-tumor tissues from the Ramazzini Institute pathology archives, arising in the Sprague Dawley rats exposed to EtOH (0% or 10%) [49] or AcA (0, 50, 250, 500, 1500 or 2500 mg/L) [50], with either chemical administered in their drinking water and supplied *ad libitum* until the spontaneous death of the animals (Fig. S1A, B). Following a comprehensive histopathological review, organs under direct exposure or metabolizing ones, which manifested increased tumor formation after EtOH or AcA treatment, were selected for downstream sequencing analysis. Tumor samples from non-exposed and exposed animals of the cheek, the Zymbal gland (a rodent auditory gland considered a direct site of exposure), the larynx, the forestomach and the liver were selected (Table S1). For each animal, unaffected kidney samples were also included as a matched germline reference. A total of twenty-eight pairs of tumor and matching normal samples from both exposure studies were ultimately included for WGS analysis (Table S1).

### Tissue processing and enrichment for tumor content

Ten sections of tumor tissue and ten sections of normal kidney of 10 μM thickness were provided for each animal from the Ramazzini Institute, Italy. The tumor sections were flanked by two additional sections stained with hematoxylin and eosin (H&E). The stained cuts were scanned using the Glissando Desktop Scanner from Objective Imaging for histopathological evaluation and delineation of the tumor area by a pathologist using Aperio eSlide Manager software (Leica Biosystems) (Fig. S2 and Table S2). For a subset of tumor samples with the actual 3D tumor size lacking (n=16), the size was based on estimates reflecting the largest tumor dimension measured on the tissue section slides (Table S2). Deparaffinization was performed by washing the slides with xylene, followed by a series of descending concentrations of ethanol as follows: two consecutive baths of xylene, followed by baths of 100% ethanol, 95% ethanol, and 70% ethanol and a final bath of distilled water. Enrichment of tumor cell content was done by macrodissection of the tumor area using a scalpel by scraping the region of interest using H&E staining of neighboring sections to assist in visualizing the area of interest for scraping. The tumor macrodissection step aimed to minimize the inclusion of non-malignant cells, such as immune cells, blood vessels, stromal cells, or necrotic tumor cells.

### DNA extraction and NGS library preparation

Genomic DNA was extracted from the fifty-six animal-matched tumor and non-tumor tissue pairs using the Macherey-Nagel Nucleospin Tissue XS kit. Ten sections of 10 μm were used for tumor tissues and three sections for non-tumor tissues. For large regions of interest (>25 mm²), a maximum of three sections was applied per column and a maximum of five sections per column for small regions of interest (<25 mm^2^). Libraries for WGS were constructed using the KAPA HyperPrep Library Preparation kit [51]. Mechanical fragmentation of DNA was performed using a Covaris S220 ultrasonicator instrument. Post-extraction DNA and final DNA libraries were quantified using the Qubit fluorometer 3.0 from ThermoFisher and an Agilent TapeStation 4200 was used to check the size distribution of genomic DNA and final libraries prior to sequencing (Fig. S3).

### Whole genome sequencing and somatic variant calling

The fifty-six libraries prepared from tumor and matched non-tumor tissues were whole-genome sequenced on the Illumina NovaSeq 6000 platform (Genewiz, Azenta) in the paired-end 150 base-pair mode (S4 flow cell), loading six to seven samples per lane to achieve ∼450 million paired-end reads per sample (Table S3). The FASTQ files were analyzed for quality using FastQC and were mapped on the rat genome reference Rnor_6.0 (Rattus norvegicus) using the BWA-MEM aligner. The somatic variant caller MuTect2 was employed to detect SBS, DBS and indels in the rat tumor samples using the matched normal kidney tissue as baseline reference for each animal to detect germline variants (Table S4). Somatic variants were filtered and annotated using ANNOVAR.

### Extraction of *de novo* mutational signatures and COSMIC signature fitting

Mutation spectra were obtained for all the tumor samples employing SigProfilerMatrixGenerator (Fig. S5). This allowed the extraction of *de novo* mutational signatures. The number of SBS signatures to be extracted was estimated based on the stability and mean sample cosine distance for different signature numbers (Fig. S6A) and four *de novo* signatures were extracted using the non-negative matrix factorization module of SigProfilerExtractor [52] (Fig. S6B and Table S5). SigProfilerExtractor was further used to decompose the *de novo* signatures into established known signatures from version 3.4 of the well curated COSMIC catalogue (Tables S6, 7).

### Attribution of activities of mutational signatures

*De novo* and COSMIC signature activities were attributed for each sample using the MSA signature attribution tool https://gitlab.com/s.senkin/MSA [53]. For COSMIC attributions, only COSMIC reference signatures that were identified in the decomposition of *de novo* signatures were included in the panel used for attribution. Optimization based on a parametric bootstrap approach was applied to extract 95% confidence intervals for each attributed mutational signature activity (Table S8).

### Analysis of mutations in cancer driver genes and pathway analysis

Somatic mutations with high variant allele frequency (VAF > 0.4), which are more often clonal events, involved in the tumor initiation and present in most tumor cells, and as a result more likely to be exposure-driven, were annotated using ANNOVAR from MAF files, and subsequently filtered to retain exonic and splicing variants with predicted functional consequences (e.g., nonsynonymous SNVs, stopgain/stoploss mutations, and indels). Gene symbols were parsed to account for multiple annotations per variant.

Two curated driver gene sets were used for annotation: IntOGen and the COSMIC Cancer Gene Census, both converted to the *Rattus norvegicus* orthologs using the babelgene R package (version 22.9) [54]. Only genes successfully mapped during ortholog conversion were retained, resulting in 616 genes from IntOGen and 724 from COSMIC. For biological processes/pathway analysis, the NIH-DAVID tool was used and only the genes mutated in at least 2 samples were considered [55] (Table S9).

### Copy Number Alterations

Somatic copy number alterations (CNAs) were identified using CNVkit (version 0.9.11) [56] on WGS data, following the recommended workflow for WGS analysis. A pooled reference was constructed by combining all available normal samples, with bias correction applied for GC content and repeat-masked regions using rn6 reference genome (Rattus norvegicus), downloaded from the UCSC Genome Browser and divided into bins of 1,000 base pairs. Gene annotation was performed using the rn6 RefFlat file, also retrieved from the UCSC Table Browser (hgTables).

## RESULTS

An elevated overall number of malignant tumors was observed in animals from all treatment groups in both the ethanol (EtOH) and acetaldehyde (AcA) exposure studies, compared to the untreated control animals (Fig. S1A, B). While the control group exhibited a background level of tumors consistent with spontaneous and sporadic tumor formation, a phenomenon commonly expected in the course of long-term rodent carcinogenicity bioassays, the treated groups demonstrated a markedly higher incidence of neoplastic lesions. This increase suggests that both EtOH and AcA exposures may promote or accelerate tumorigenesis beyond the baseline levels observed in untreated animals, potentially through mechanisms involving cellular stress, DNA damage, or other carcinogenic pathways.

We observed a statistically significant trend toward earlier spontaneous death in the treated animals, which on average died around 104 weeks of age, compared to 129 weeks in the untreated control group, considering all analyzed tumor sites. This difference, similarly observed upon separate analysis for head and neck tumors, may reflect accelerated tumor development following exposure to ethanol (EtOH) or acetaldehyde (AcA), ultimately impacting animal survival (Fig. 1A–D). Furthermore, tumors were significantly larger in treated animals than comparable tumors arising spontaneously in untreated controls (Fig. 1E–H). The effects of treatment on both survival and tumor size were particularly pronounced in head and neck tumors —affecting sites such as the cheek, Zymbal gland, and larynx— (Fig. 1C, D, G, H), and these differences were further accentuated when EtOH- and AcA-treated groups were analyzed together (Fig. 1C, G).

**Figure 1.**
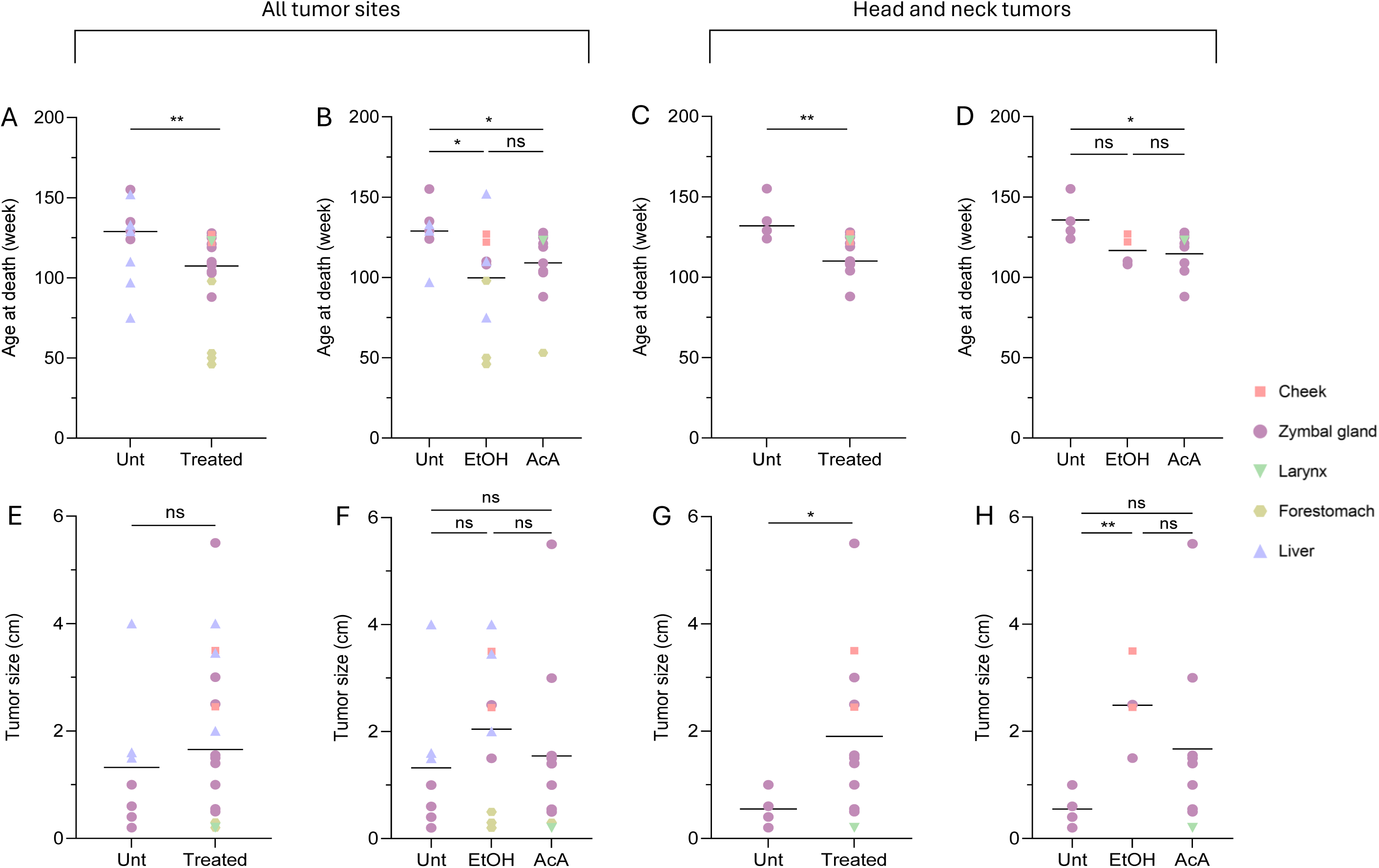
The age-at-death and tumor-size effects of EtOH- or AcA exposures in Sprague Dawley rats. (**A-D**) The graphs show the age of the untreated control and exposed rats categorized by cancer site, at the time of spontaneous death. (**E-H**) Graphs show the size of the tumor samples, categorized by cancer site. All tumor sites (**A, B, E, F**) or a subset of head and neck tumors (**C, D, G, H**) are shown separately. Untreated and treated samples (combining EtOH- and AcA-treated samples) are shown in (A, C, E, G). Untreated and the samples grouped by individual EtOH and AcA exposures are shown in (B, D, F, H). Unt: Untreated, Treated: EtOH- or AcA-treated combined. * *p*<0.05, ** *p*<0.01 using non-parametric Mann-Whitney test (combined EtOH and AcA treatment groups) or Kruskal-Wallis test (separate treatment groups) followed by uncorrected Dunn’s test for multi-group comparisons. n.s.: non-significant *p* value. Median is represented by black horizontal lines.

We analyzed overall somatic mutation frequency in whole-genome sequenced tumor samples and found that somatic mutation counts, specifically single-base substitutions (SBS), were relatively similar across tumors from control, EtOH-treated, and AcA-treated animals (Fig. S4A). Three samples displayed markedly elevated mutation numbers, potentially indicating a hypermutator phenotype or technical artifacts related to sample quality, DNA preparation, or sequencing. The majority of mutations were SBS events, and they were similarly distributed across the genome in all tumor sites and exposure groups, with most located in intergenic and intronic regions, as expected (Fig. S4B, C).

### Exposure-related enrichment of signature SBS17

We then explored the SBS-based mutational signatures in the rat tumors. We identified four *de novo* SBS mutational signatures (SBS96 A, B, C, D) of high stability (Fig. S6) and subsequently decomposed them into COSMIC mutational signatures (Fig. 2A, Fig. S7 and Table S6). The relative per-tumor distribution of the decomposed COSMIC SBS signatures obtained using SigProfilerExtractor detected the clock-like SBS1, SBS5 and SBS40a signatures present across all or most tumors, regardless the treatment condition. We also observed the sporadic, low-representation occurrence of signatures SBS30 and SBS37, with no specific association with treatment conditions. Interestingly, signatures SBS17a and SBS17b were enriched in the treated head and neck tumors, including one cheek (CH_8) and 27% of the Zymbal gland tumors (ZG_9, ZG_23, ZG_24, ZG_25) derived from animals exposed to 10% EtOH and higher doses of AcA (1500 and 2500 mg/L) (Fig. 2B). An optimized mutational signature attribution approach by using MSA [53] confirmed the extraction/decomposition results mentioned above (Fig. 2C) and the enrichment the same five samples of both SBS17b and SBS17a (Fig. 2D and Fig. S8, respectively). While the signal for the T>G-based SBS17b appeared highly specific and validated the SigProfilerExtractor findings, with MSA we observed some lower-level attribution interference for the T>A, T>C-based SBS17a in a subset of samples. The overall quantitative summary of the detection of SBS17a/b in treated *vs* untreated head and neck tumors is shown in Fig. 2E, F. Apart from the SBS signatures described above, we detected three COSMIC mutational signatures SBS45, SBS58 and SBS95 that are considered possible technical artefacts (Fig. 2B, C and Fig S7). These patterns were extracted from the data as SBS96C *de novo* signature which is artefact-derived for approximately 90%. These artefactual signatures were observed somewhat more frequently in tissues from EtOH-treated rats, suggesting study-specific differences in sample processing or quality. Lastly, the analysis of single base substitution (SBS) signatures in the tumor tissues revealed no evidence of SBS16 formation, a mutational signature previously tentatively associated with alcohol consumption in humans. This absence suggests that in the tumors of EtOH- and AcA-treated rats, these compounds do not elicit this specific mutational pattern.

**Figure 2.**
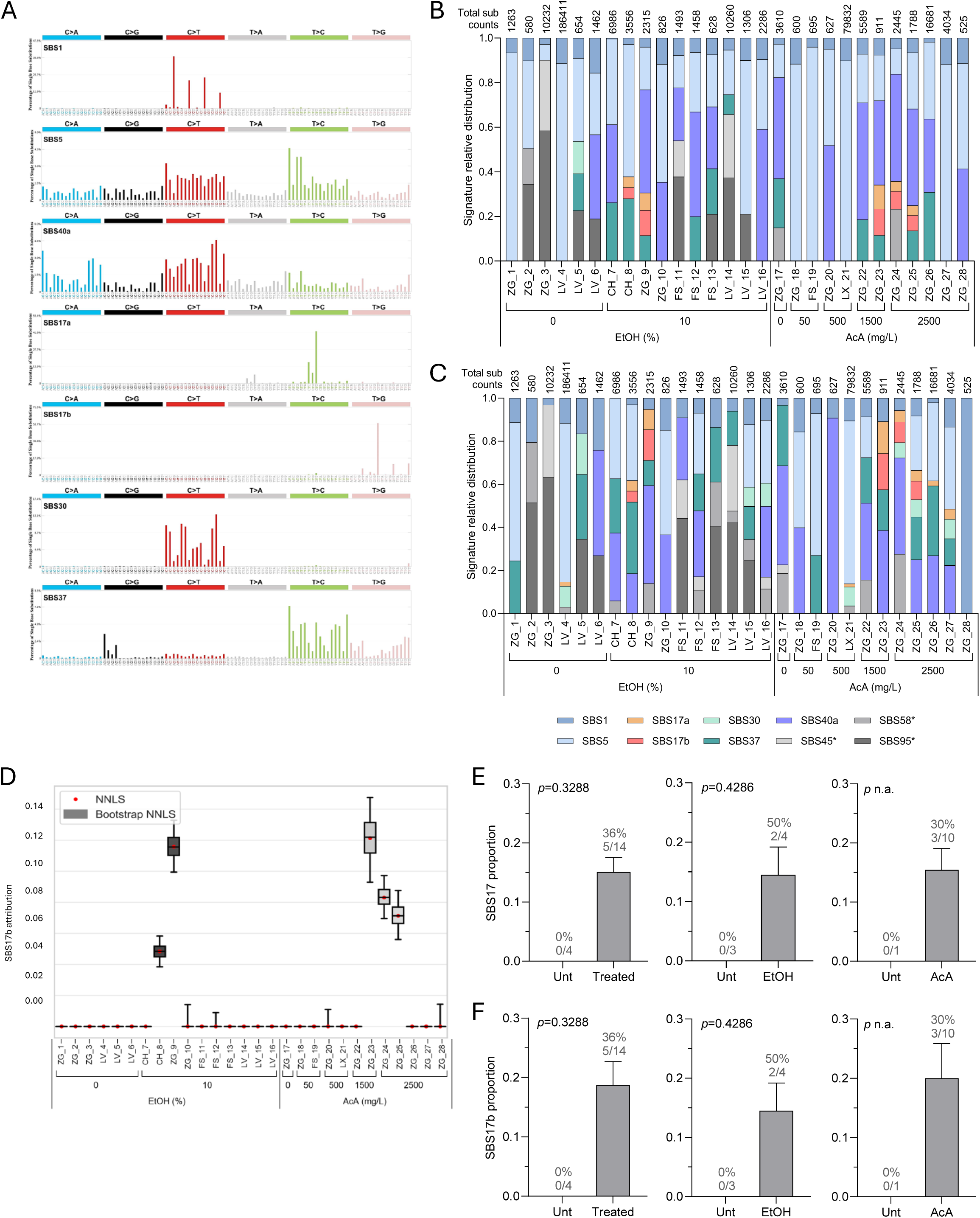
SBS mutational signatures identified in the rat tumors. (**A**) COSMIC SBS mutational signatures of biological (non-artifact) origin extracted from the sequenced tumor samples. (**B, C**) Graphs show relative distribution of the mutational signatures across tumor samples with related total mutation counts on the top. * = signatures listed in COSMIC as possible sequencing artifacts. Mutational signatures were extracted either by using SigProfilerExtractor (B) or attributed by using Mutational Signature Attribution (MSA) (C). (**D**) Relative attribution of SBS17b across tumor samples by using optimized attribution by the MSA tool. (**E**) Summary of the presence of signature SBS17 (SBS17a and SBS17b combined) in head and neck tumors (cheek, Zymbal gland, larynx) of animals exposed to either treatments (left, EtOH and AcA combined), EtOH only (middle) or AcA only (right) using SigProfilerExtractor. (**F**). Validation of the presence of signature SBS17 (considering only SBS17b) in head and neck tumors (cheek, Zymbal gland, larynx) of animals exposed to either treatments (left, EtOH and AcA combined), EtOH only (middle) or AcA only (right) using MSA. Statistical analysis done using nonparametric Mann-Whitney test. n.a.: *p*-value non applicable. Abbreviations: NNLS - non-negative least squares.

### The identified DBS and indel signatures do not trail with exposure

To further investigate the mutational patterns in the tumors of the animals, we analyzed the double base substitution (DBS) and small insertion and deletion (indel or ID) signatures (Table S4). Two *de novo* DBS signatures (and five *de novo* ID signatures were extracted in the tumor samples and decomposed into reference COSMIC signatures (Fig. S9, S10 and Tables S5, S6). However, we did not observe clear exposure-related differences related to DBS and indel signature distribution. Similarly to SBS16, signatures DSB4 and ID11 previously tentatively associated with the combination of alcohol drinking and smoking [33] were not detected in the EtOH- or AcA-treated rat tumor genomes.

### Exposure-related profiles of cancer driver gene mutation

We next explored the potential cancer driver mutations and their distribution across the treatment conditions. Among the candidate cancer driver genes analyzed, several well-established genes with pivotal roles in oncogenesis were detected as non-silently mutated in the tumors (Fig. 3). Analysis of non-silent mutations in cancer driver genes highlighted a noteworthy divergence between untreated and EtOH/AcA-induced tumors, with treated samples harboring recurrent mutations in well-established driver genes. These included the tumor suppressor gene Tumor protein p53 (*Tp53*), harboring established or likely loss-of-function amino acid substitutions (p.G106A, p.T123M, p.Y124C, p.A157V, p.V141A, p.S239F, and p.G264R), which are likely to disrupt p53’s tumor-suppressive function. Additionally, the gene mechanistic target of rapamycin (*Mtor*) carried recurrent mutations (p.R731H and p.A1344D), though the functional significance of these variants remains unknown. These alterations were found solely or primarily in tumors from treated animals. In contrast, the gene Tenascin C (*Tnc*), which encodes a large extracellular matrix glycoprotein, was mutated exclusively in tumors from untreated animals, featuring a possibly deleterious substitution (p.R1455C) alongside variants of uncertain effect (p.Q1554Q, p.T1273I, p.L1722F, and p.A1863T). Lastly, independently of the exposure condition, we observed driver mutations in the oncogene Harvey rat sarcoma viral oncogene homolog (*Hras*), with gain-of-function substitutions (p.G12E and p.Q61L), both well-established oncogenic driver mutations associated with aggressive, proliferative tumors in many human cancers. Nevertheless, the overall mutation rates in cancer driver genes were low, consistent with the weak mutagenicity in rats as suggested by the relatively low mutation burden and lack of pronounced treatment-associated mutational patterns.

**Figure 3.**
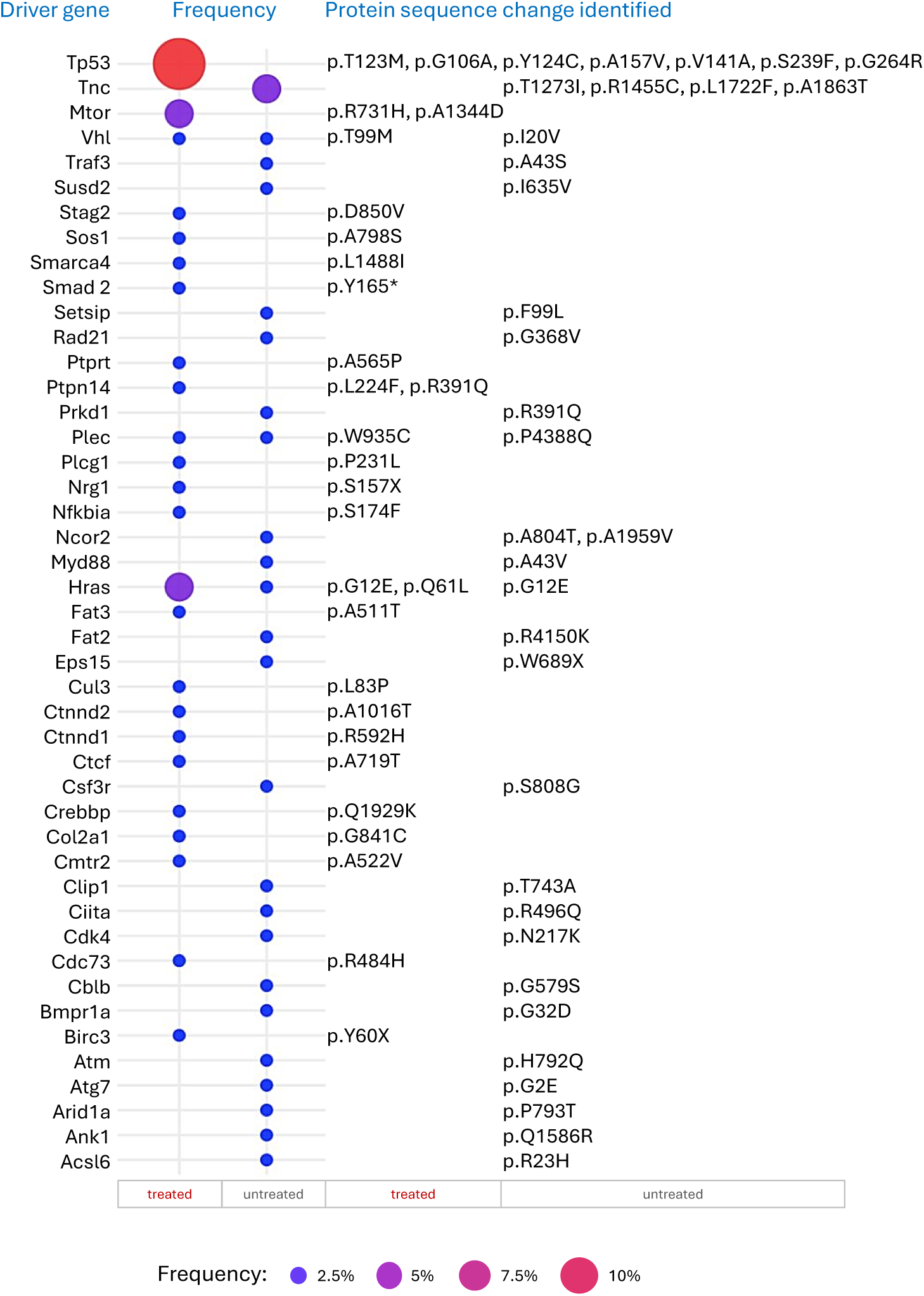
Identification of the mutated cancer driver genes. The graph shows the distribution of the mutated cancer driver genes across treated and untreated sample groups. The relative frequency of the number of samples affected by at least one mutation occurring in the listed cancer driver genes are represented by the colored dots. The corresponding changes in protein (amino acid) sequence are detailed on the right of the graph. Refer to Methods for additional details.

### Copy number alterations are not associated with EtOH/AcA treatment

Comprehensive analysis of copy number alterations (CNAs) across the whole-genome sequencing data did not reveal any recurrent or well-established amplifications or deletions in the analyzed tumor samples. The observed copy ratio profiles remained largely within the expected diploid range, with only isolated and low-amplitude fluctuations not reflecting the treatment conditions (Fig. S11). These minor variations appeared to be random and did not correspond to any known genomic hotspots or regions typically affected by therapy-induced genomic instability. Overall, the CNA landscape suggests a stable genomic architecture in response to the treatment, with no evidence of widespread genomic gains or losses that might indicate selective pressures or clonal evolution under either of the exposure regimens. These findings indicate that large-scale genomic instability, as manifested through CNAs, is not a prominent feature underlying tumor development in animals treated with either EtOH or AcA.

## DISCUSSION

Chronic treatment of rats with either EtOH or AcA enhanced the formation of tumors compared to untreated control animals, with pronounced effects on survival and tumor size, especially in the context of head and neck cancers. Genomic analysis of mutational patterns formed in those tumors revealed an exposure-specific enrichment in signature SBS17 formation in a subset of head and neck tumors from ethanol- and high dose AcA-exposed animals, specifically cheek and Zymbal gland. We propose an association between the formation of SBS17 and the chronic consumption of alcohol. Interestingly, most of the organs where SBS17 has been found enriched are also cancer sites associated with alcohol drinking (namely esophagus, breast, stomach, pancreas, colorectum) [18], suggesting that the signature formation may be a consequence of EtOH or AcA exposure upon alcohol consumption (Fig. 4A).

**Figure 4.**
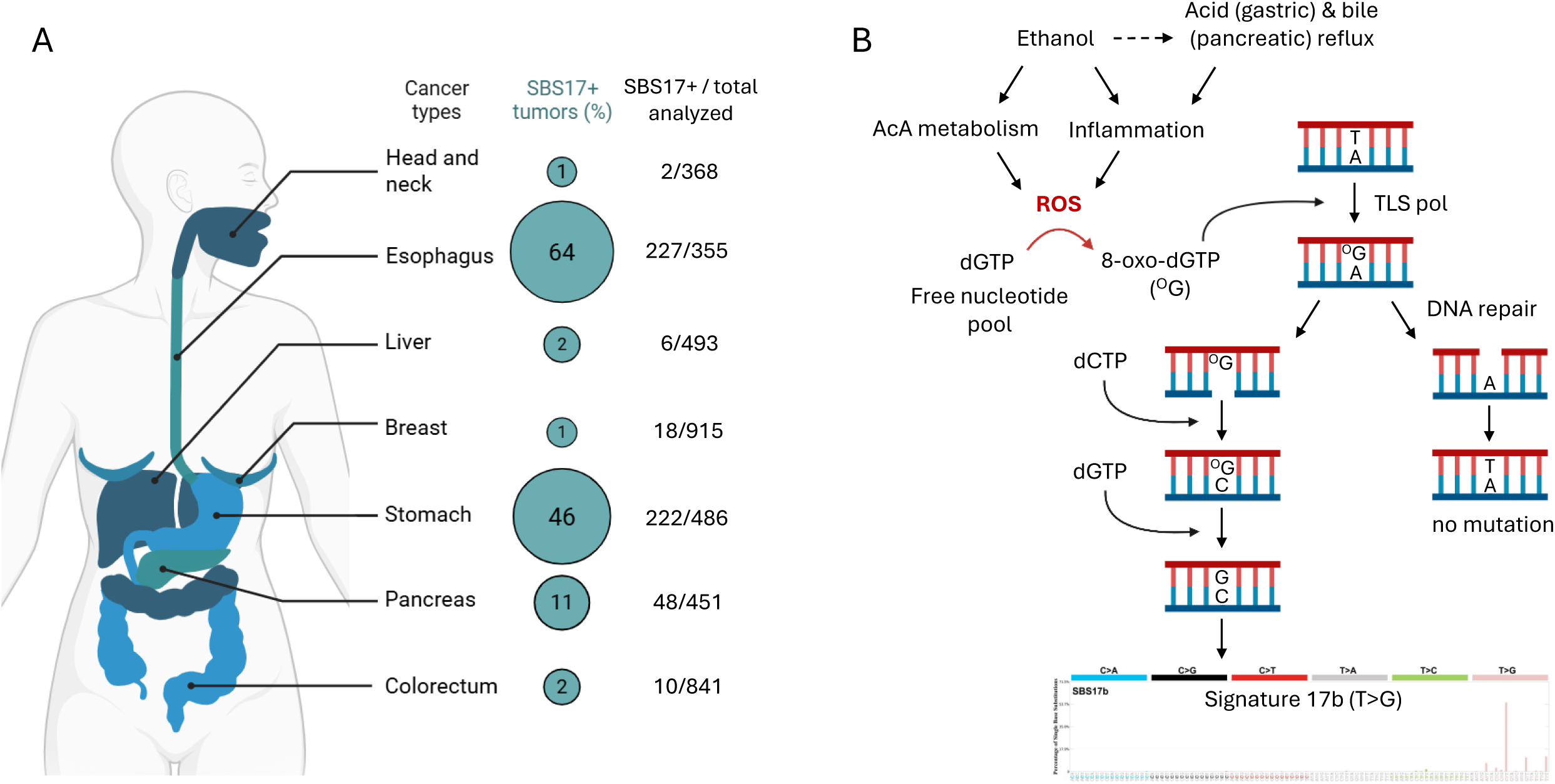
Putative model associating alcohol exposure and SBS17 formation. (**A**) The diagram shows alcohol-associated cancers and their relative enrichment for signature SBS17, shown as % of tumors positive for SBS17 among the analyzed tumors of the same type (source: COSMIC). The proportion of positive samples and total of analyzed tumors are shown on the right. (**B**) Proposed mechanisms by which oxidative stress (reactive oxygen species, ROS) can be induced by ethanol use through AcA metabolism and/or chronic inflammation, also possibly triggered by acidic reflux (gastric or pancreatic). ROS can directly affect the free nucleotide pool and oxidize dGTP to 8-oxo-dGTP which may in turn induce T>G transversions via mispairing in DNA with dA and ultimately lead to SBS17b-like T>G-enriched mutation pattern.

To date, the exact mechanisms driving the formation of signature SBS17 are not fully characterized, although several hypotheses have been proposed, the main one being the implication of oxidative DNA damage in the formation of mutation patterns specific for SBS17 [57–59]. The development of Barrett’s esophagus and its transformation to esophageal adenocarcinoma, where SBS17 is frequently observed, is mostly associated with chronic gastro-esophageal reflux (acids), responsible for persistent inflammation and accumulation of reactive oxygen species (ROS) resulting in greater oxidative stress and related DNA damage [60–63].

Pancreatic reflux containing bile and pancreatic enzymes also affects the stomach and esophagus in the most severe cases. Besides damaging the DNA-incorporated bases, acid and bile-derived oxidative stress can also directly affect the free nucleotide pool, and form modified bases such as 8-oxo-dGTP from oxidation of dGTP. Incorporation of 8-oxo-dGTP into DNA by translesion synthesis (TLS) polymerases subsequently interferes with DNA replication and repair pathways and increases the risk of T>G transversions, the hallmark of SBS17 [59, 64, 65] (Fig. 4B). In keeping with the proposed role for inflammatory processes in inducing oxidative stress and SBS17, reduced frequency of the signature 17-specific point mutations has been reported in patients with Barrett’s esophagus who used non-steroidal anti-inflammatory drugs [66]. In human cancers, SBS17 is often observed in gastrointestinal tumors, such as esophageal adenocarcinomas [67, 68], its precursor Barrett’s esophagus [60, 69], and in gastric cancers [70], where chronic inflammation and gastric and pancreatic reflux of acids and bile contribute to persistent oxidative stress. Importantly, chronic alcohol consumption can be a risk factor for developing gastroesophageal reflux disease [71–73], and is a known inducer of ROS via ethanol/AcA metabolism, and may exacerbate oxidative DNA damage and ultimately trigger mutational processes linked to SBS17.

However, the formation of SBS17 is not exclusive to alcohol exposure. Multiple studies have reported its presence in rodent tissues in the absence of any known exogenous carcinogen. In cultured mouse embryonic fibroblasts (MEF), the signature formed naturally in cultures undergoing senescence-bypass and clonal-immortalization [15, 74–76]. In addition, SBS17 arose in subclones of cell populations expanded from mouse bowel single organoid stem cells under prolonged culture conditions [77] and in murine neonatal single-cell fibroblasts [78]. Formation of SBS17 was also reported in *in vivo* rodent models. In mice treated with aflatoxin B1 that developed liver HCC-like cancer, SBS17 arises as a late onset pattern, suggesting its formation is independent from the experimental exposure [46]. Further, mouse models of diffuse human gastric cancer show a mutational pattern that is reproducible with human gastric cancer, including the presence of SBS17 [79]. Genome-wide analysis of colorectal tissues and the comparison of somatic mutations across 16 different species shows the presence of SBS17 only in rodent cases (mouse, rat, naked mole-rat) [80]. Finally, the mutational pattern was similarly found in mouse tissues from both tumor and normal samples [47]. These observations suggest that SBS17 may, at least in part, reflect endogenous DNA damage processes exacerbated under certain stress conditions such as those induced by exposure to alcohol and/or oxidative stress. On another hand, the relatively low amount of overall mutational events observed in tumor samples collected from exposed animals, and the lack of a strong association between one or more mutational signatures with the treatment, proposes that direct mutagenicity may not be the most prominent mechanism for alcohol-induced carcinogenesis. Although ethanol EtOH and AcA have traditionally been classified as genotoxic and mutagenic agents, growing evidence suggests that mutagenicity may not be the predominant mechanism through which they drive carcinogenesis. While AcA is known to form covalent DNA adducts such as N²-ethyl-2’-deoxyguanosine (*N*²-Et-dG) and induce DNA-protein and interstrand crosslinks [5, 81, 82], these lesions do not necessarily translate into permanent mutations. Many AcA-induced modifications are bulky, transient, or effectively repaired before DNA replication, thereby evading detection by standard NGS approaches as applied in this study, which primarily identify stable DNA sequence alterations [81]. For example, interstrand crosslinks may stall DNA polymerases and activate repair pathways such as the Fanconi anemia pathway without causing detectable sequence changes [83, 84]. Similarly, DNA adducts like *N*²-Et-dG are relatively unstable and may be removed under physiological conditions, further questioning their direct mutagenic impact *in vivo* [5]. Moreover, the mutagenic potential of EtOH and AcA may be context-dependent and secondary to other cellular stress responses [85, 86]. AcA may promote carcinogenesis through non-mutagenic mechanisms, such as epigenetic alterations. It has been demonstrated that AcA can disrupt histone modification patterns and DNA methylation, potentially leading to transcriptional dysregulation without direct DNA sequence changes [87, 88]. Chronic alcohol consumption can also contribute to the emergence of an inflammatory microenvironment or other types of microenvironment changes leading to a more permissive setting, allowing the selection, growth and spreading of pre-cancerous cells [89]. Importantly, EtOH and AcA metabolism pathways are closely linked to the induction of oxidative stress, a key mechanism underlying alcohol-related carcinogenesis [86, 90, 91]. As mentioned previously, SBS17 formation may be the consequence of ROS accumulation, therefore alcohol-induced oxidative stress may contribute to the formation of SBS17 in the context of chronic alcohol consumption. These findings suggest that the role of EtOH and AcA in cancer development may involve a complex interplay of genotoxic and non-genotoxic mechanisms, and that mutagenicity may not be the sole or even primary driver of pathogenicity. We speculate that the consequences of EtOH/AcA exposure resulting from alcohol drinking are probably multiple, leading to a combination of deleterious effects at a cellular level, but also at a broader level changing the microenvironment properties, that would not necessarily be detectable through genomic analysis using whole genome sequencing.

Our study presents a considerable technical innovation; the analyzed animal samples were alcohol-fixed paraffin-embedded (AFPE) which differs from the formalin-fixed tissues (FFPE), most commonly used for WGS analyses [92, 93]. Alcohol fixatives have been shown to better preserve tissue morphology, protein, and nucleic acids than the formalin-based fixation [94]. This possibly ensures higher quality sequencing data with reduced sequencing artefacts related to fixation and tissue processing, ultimately improving the extraction of exposure-associated mutational signatures in comparison to results from recent multiple FFPE-based studies, including our own [93–96].

This study has some limitations, primarily due to the relatively small number of tumor samples available, which restricts the statistical power of the analysis. In addition, tumors from different organs were combined, although the direct effects of EtOH and AcA exposure may vary depending on the organ. It is therefore conceivable that multiple mechanisms are involved in the response to EtOH/AcA exposure, depending on the tumor site. Despite these limitations, however, our study represents an important advance, as it is the first of its kind to investigate the much-debated formation of alcohol exposure-specific mutational signatures or higher-level genomic DNA alterations, using a mammalian *in vivo* cancer model and as such provides novel insights into the potential mechanisms of alcohol-related carcinogenesis.

## CONCLUSION

Taken together, our findings suggest a potential relationship between alcohol consumption and the formation of SBS17, implicating the accumulation of oxidative damage and stress induced by ethanol metabolism pathways and other indirect stress triggered by the exposure. The partial presence of SBS17 in exposed animals, coupled with the absence of presumed alcohol-associated signatures (e.g., SBS16), supports the notion that in the context of *in vivo* bioassays in rats neither ethanol nor AcA exposure robustly induce specific and consistent sequence-altering mutations detectable by NGS. We propose that the formation of SBS17 itself, likely due to oxidation of the free nucleotide pool, may become exacerbated in rodents due to exposure related-stress and the weaker repair mechanisms in rodents compared to humans, but is not directly attributable to direct mutagenic effects of AcA.

Other mutation types, including DBS, indels, and CNAs, did not indicate consistent changes associated with EtOH or AcA exposure. Analysis of cancer driver genes revealed few recurrent mutations, albeit with low mutation frequencies across tumors. Some known drivers such as *Tp53* and *Mtor* were recurrently mutated in exposed animals, including established hotspot mutations, but the overall low mutation rates support the idea that direct mutagenicity is not the primary mechanism of alcohol-driven carcinogenesis in this model.

Unrepaired lesions can lead to replication errors and genomic instability, resulting in mutations and chromosomal aberrations. The lack of exposure-related strong mutational signatures observed in this study does not imply the absence of genotoxic stress. Instead, it underscores the limitations of mutation-centric detection methods and the need for complementary analytical strategies, such as DNA adductomics, crosslink-specific assays, analyses of the epigenome or microenvironment changes, among others to fully characterize the biological impact of EtOH/AcA exposure. Collectively, these findings suggest a more nuanced model in which non-mutagenic mechanisms may play a central role in alcohol-associated carcinogenesis, despite the mutagenic potential of AcA-derived DNA damage.

## DATA AVAILABILITY

Aligned WGS reads for the rat tumors from this study have been submitted to the NCBI BioProject database under accession number **PRJNA1282587**. For individual BioSample accession numbers, refer to **Supplementary Table S10**.

## AUTHOR CONTRIBUTIONS

BC carried out experiments and generated the main data. ET and DM provided animal samples and ET and BAA conducted histopathological evaluation of the tumor samples. CC processed the archived tissue samples for DNA extraction and library preparation performed by BC. Analysis of the data was carried out by BC, VC, SS, FV, JZ. Key aspects of the study were overseen by MK, ZH, SB, DM and JZ. The manuscript was written by BC and JZ. All authors reviewed the results and revised the manuscript. All authors approved the final version of the manuscript.

## SUPPLEMENTARY FIGURE LEGENDS

**Supplementary Figure S1.**
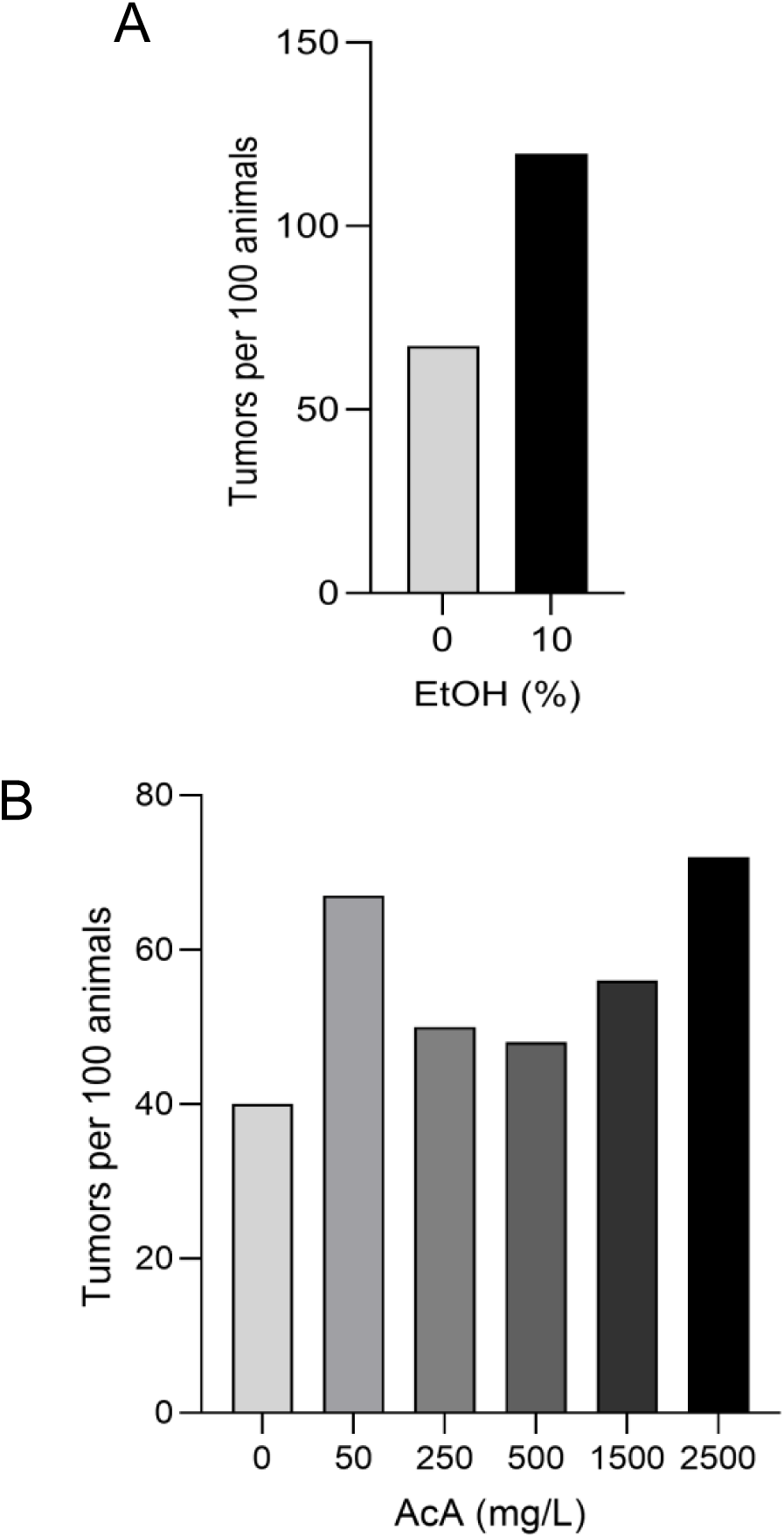
Effects of EtOH or AcA on malignant tumor formation in Sprague Dawley rats. (**A-B**) Tumor burden per 100 animals in (A) the EtOH treatment group and (B) the AcA treatment group, with doses indicated.

**Supplementary Figure S2.**
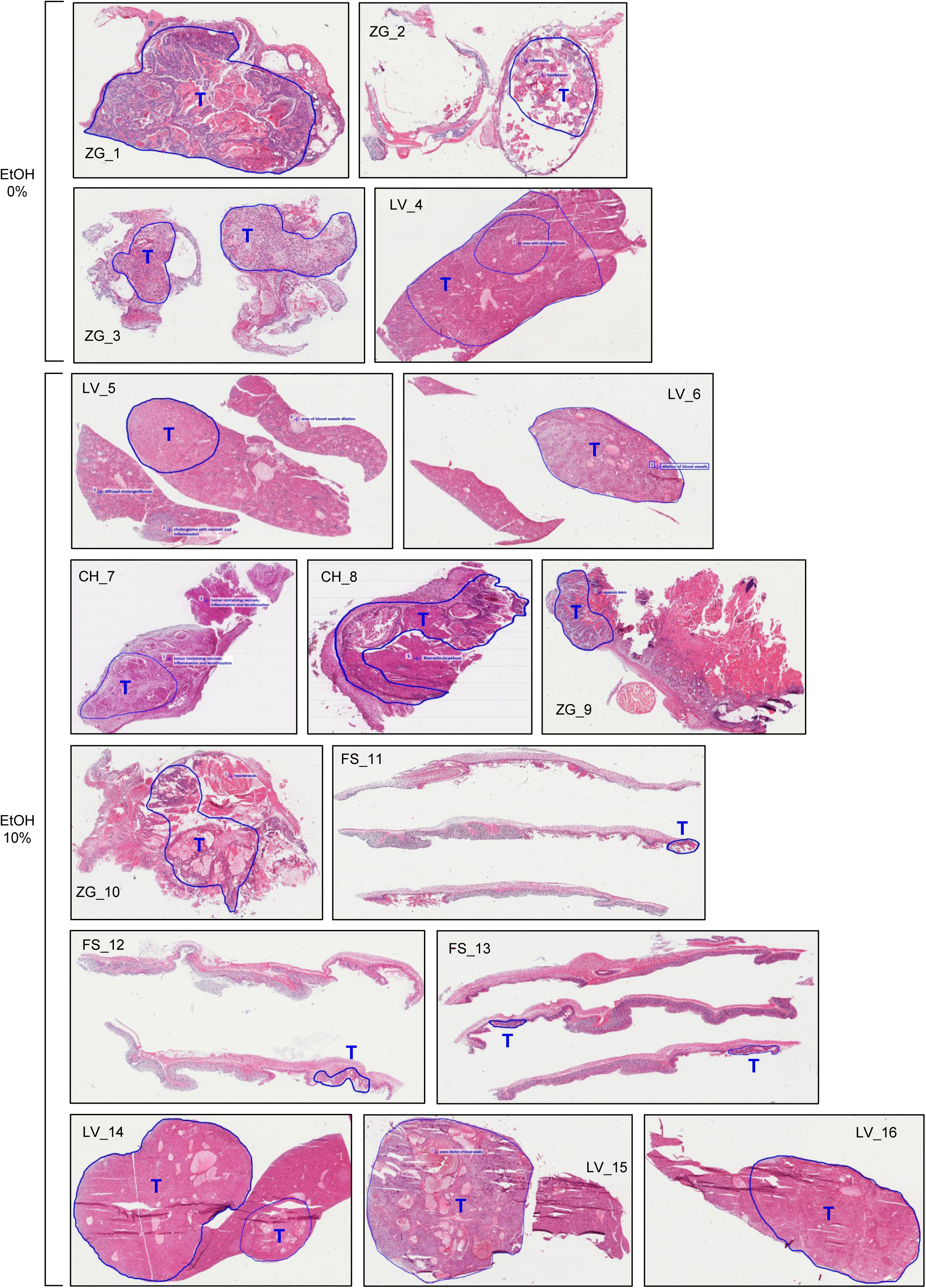
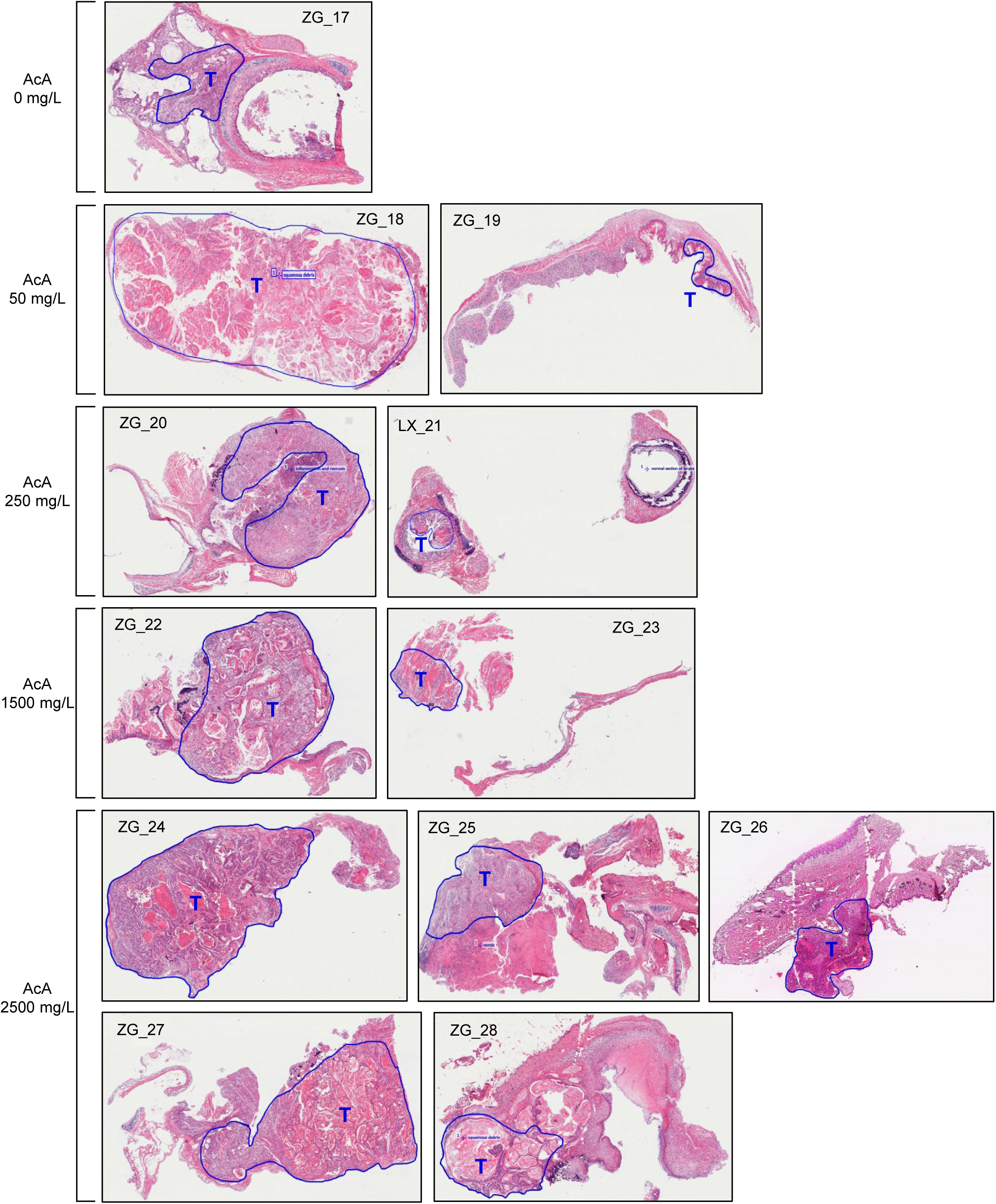
Demarkation of tumor mass areas on hematoxylin eosin-stained scans of the rat tissue samples. Scans of the 28 tumor tissue sections stained with hematoxylin and eosin and their annotation for tumor area (T, in blue), for manual macrodissection of tumor-enriched material. No scans of the animal-matched non-tumor tissues are shown as the entire sections of the control tissues were used for DNA preparation. Exposure groups are indicated on the left side. Abbreviations: AcA - acetaldehyde, EtOH - ethanol, CH - cheek, ZG - Zymbal gland, LV - liver, FS - forestomach, LX - larynx, T - tumor area.

**Supplementary Figure S3.**
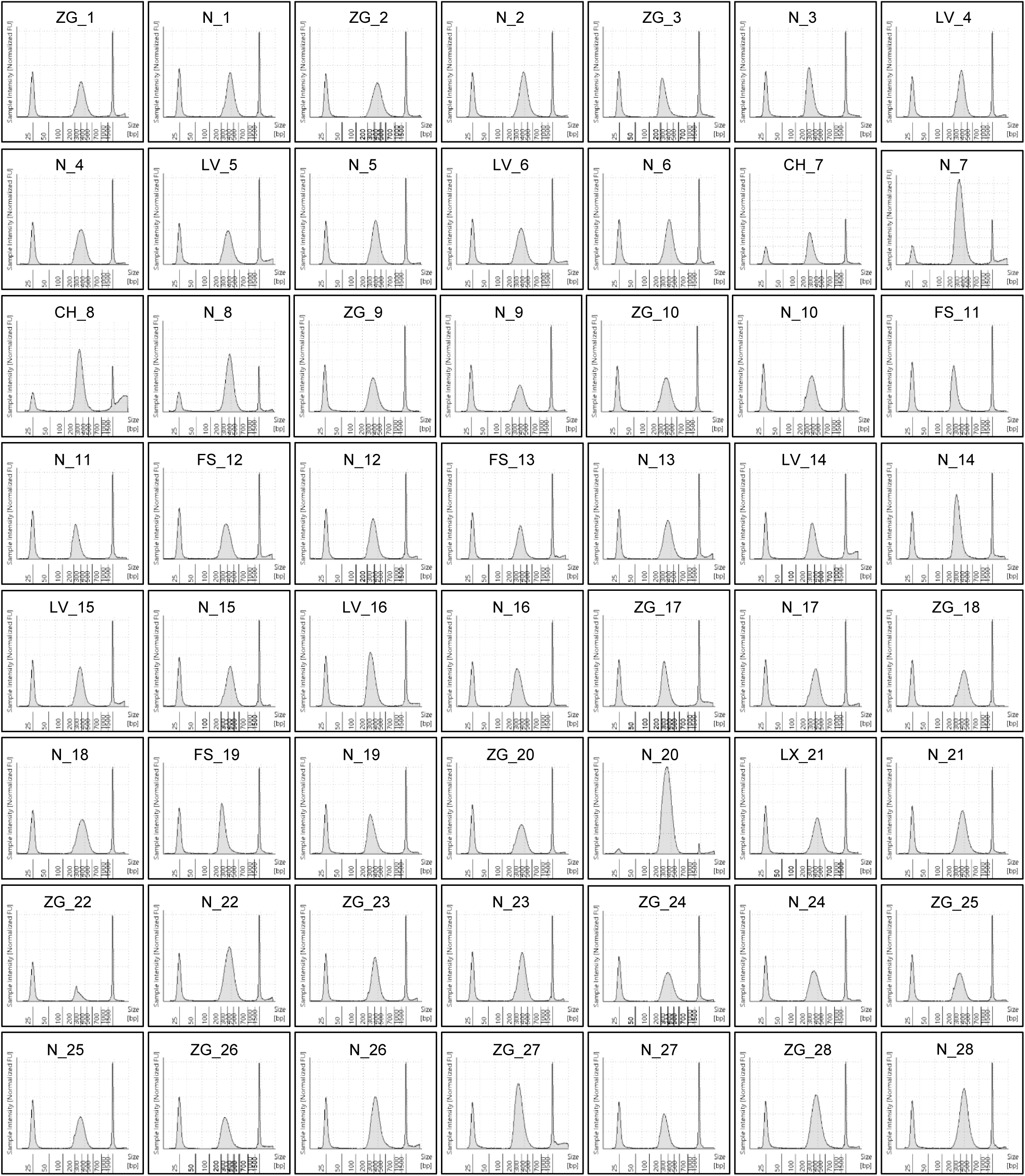
Whole genome sequencing DNA library profiles. Fragment size distribution of the 56 DNA libraries for WGS prepared from the rat tumor and normal samples is shown. The libraries were analyzed for quality and uniformity by using the Agilent TapeStation 4200. CH - cheek, ZG - Zymbal gland, LV - liver, FS - forestomach, LX - larynx, T - tumor tissue, N - non-tumor tissue.

**Supplementary Figure S4.**
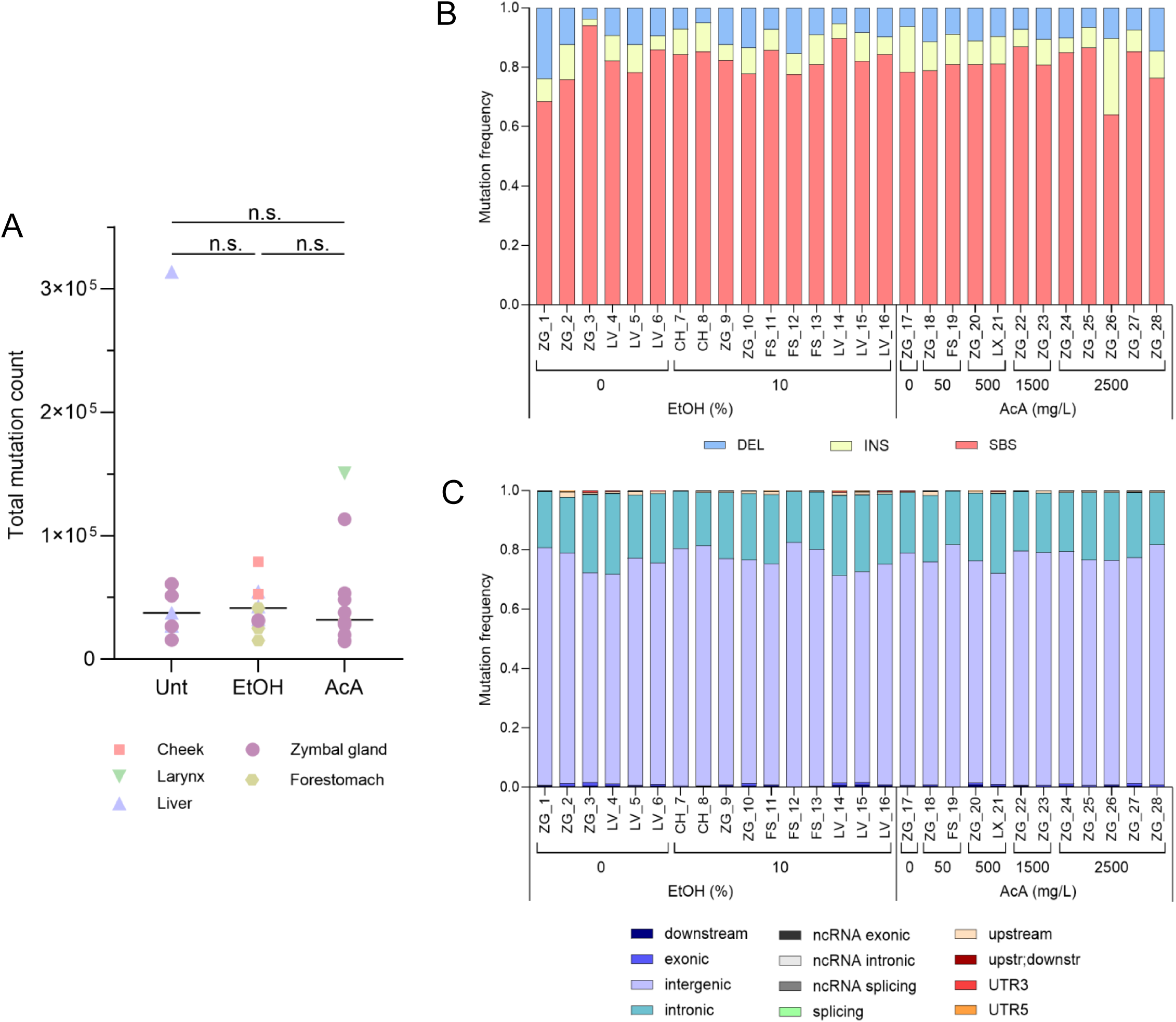
Summary of mutation burdens and types in rat tumor samples. (**A**) The graph shows the total mutation counts in tumor samples collected in the untreated, EtOH- and AcA-treated rats. Non-parametric evaluation of the entire group was performed using Kruskal-Wallis test, followed by multiple paired comparisons using uncorrected Dunn’s test; n.s. -non-significant *p*-value. (**B**) Distribution of main mutation types found in the sequenced tumor samples. DEL - deletion, INS - insertion, SBS – single nucleotide substitution. (**C**) Distribution of SBS mutations based on genomic segments. nc – non-coding, UTR – untranslated region.

**Supplementary Figure S5:**
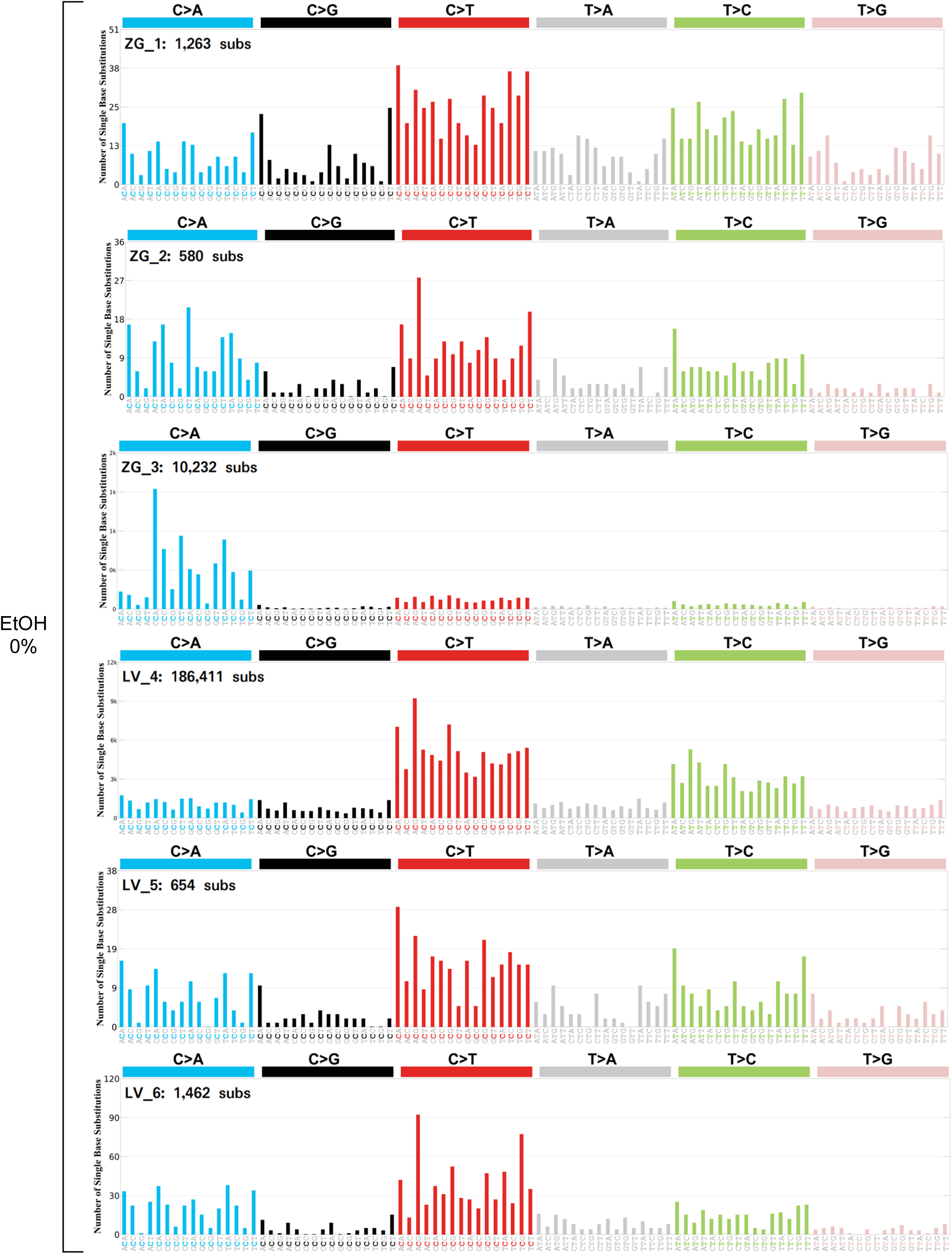
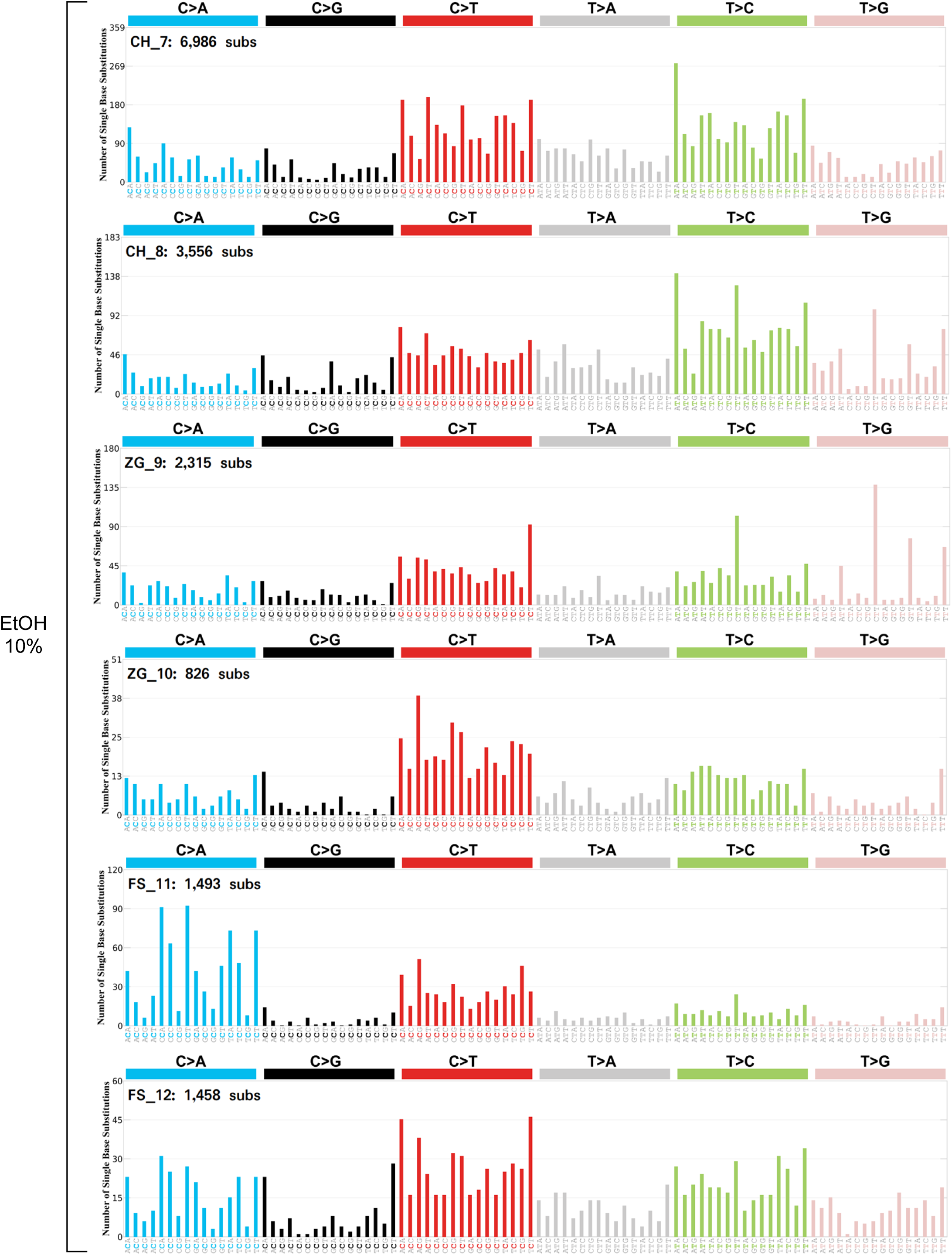
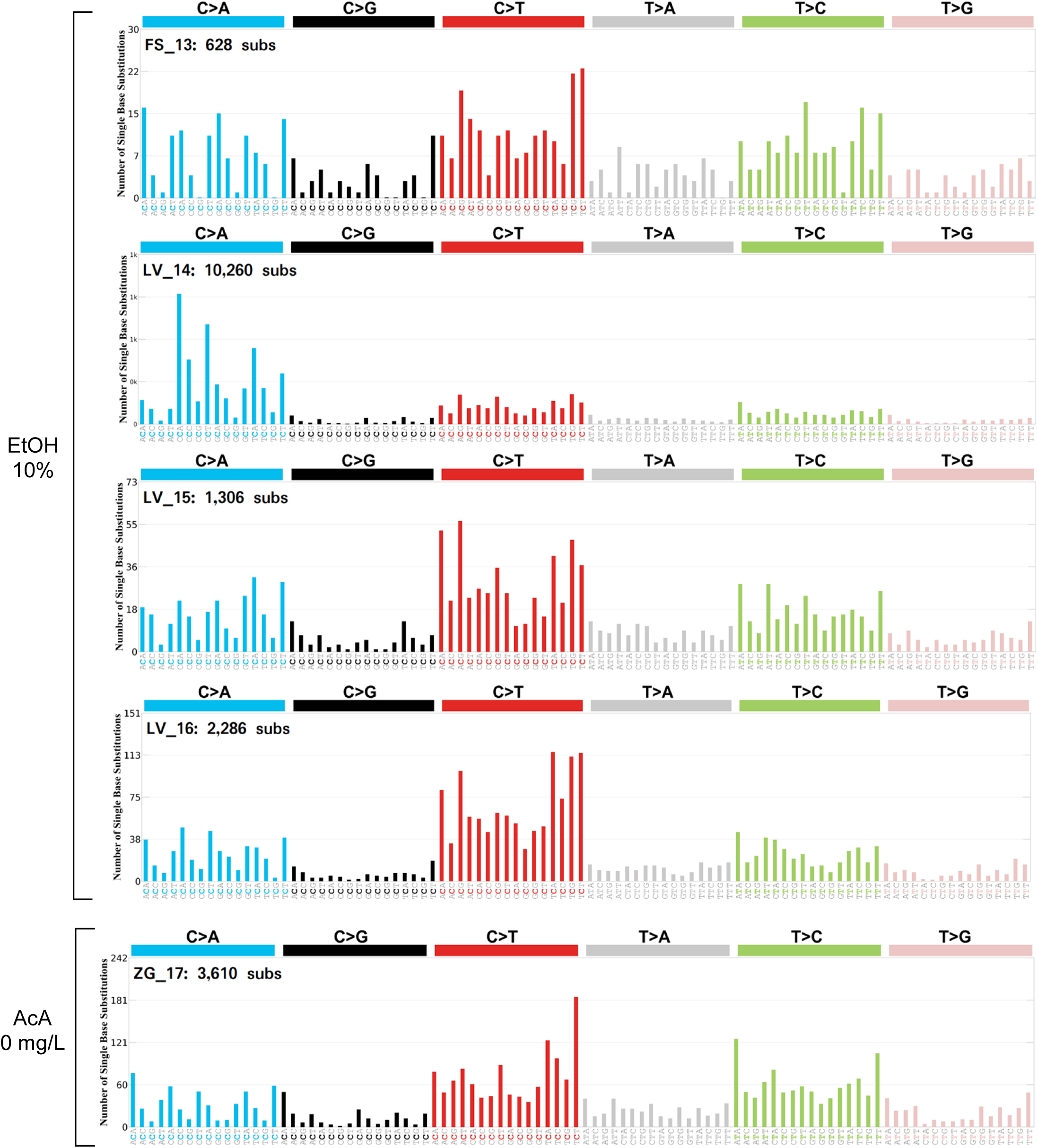
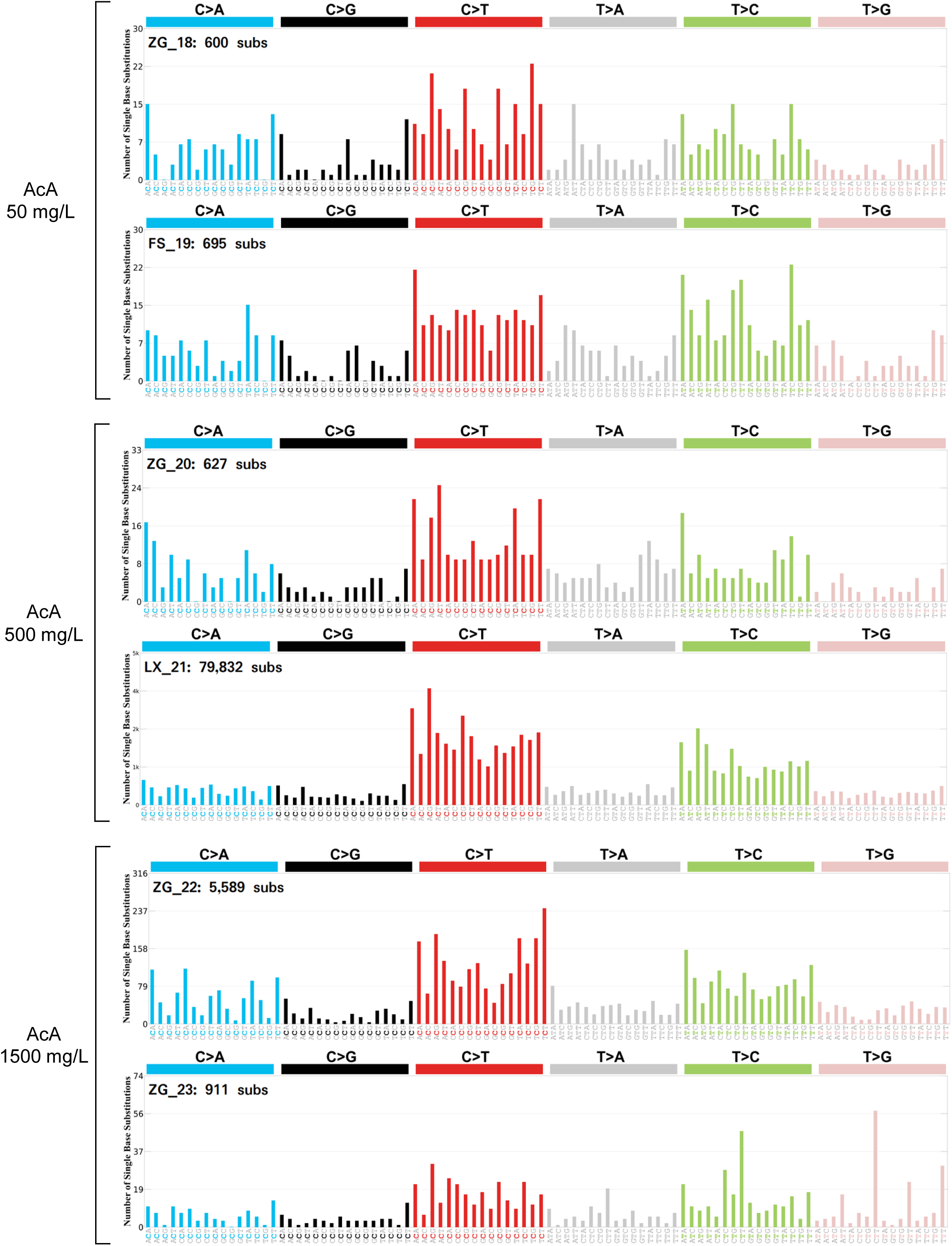
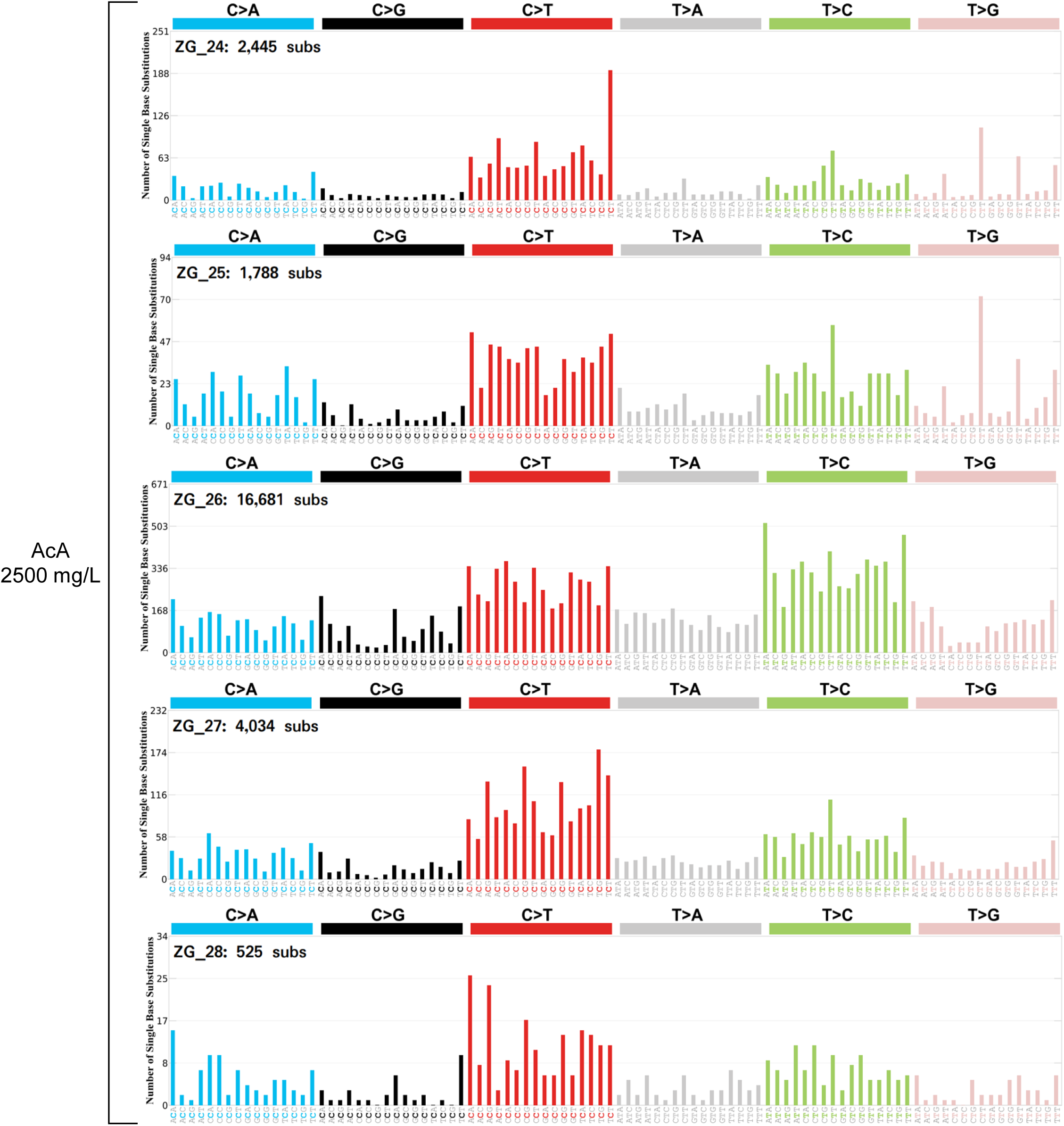
Whole-genome somatic mutation spectra in individual tumors. Histograms show trinucleotide mutation spectra (96-channel) with total substitution counts, resulting from WGS analysis of the 28 tumor samples collected from untreated animals, or animals exposed to EtOH or AcA at various doses (as indicated to the left of the histograms). Abbreviations: CH - cheek, ZG - Zymbal gland, LV - liver, FS - forestomach, LX - larynx,

**Supplementary Figure S6.**
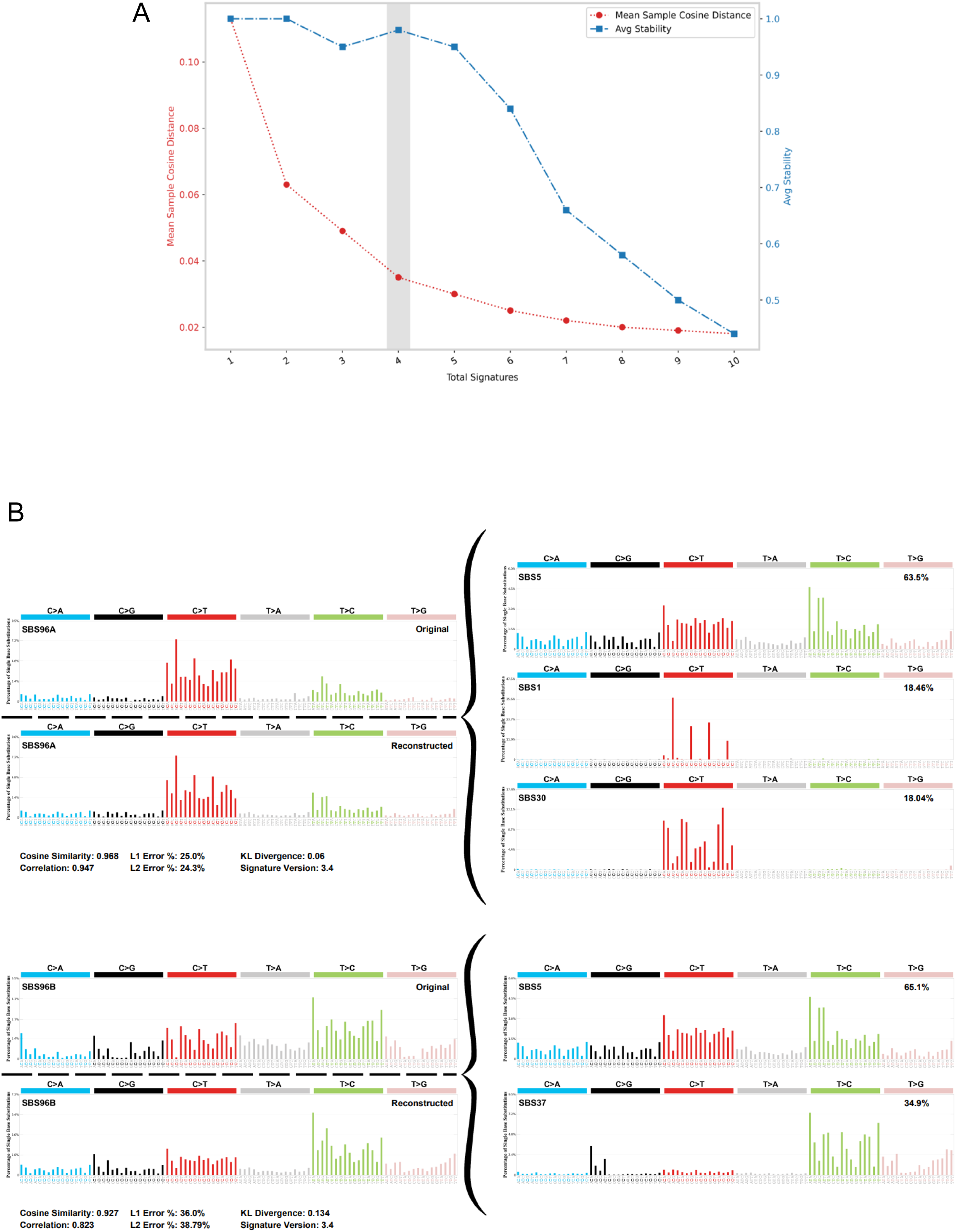
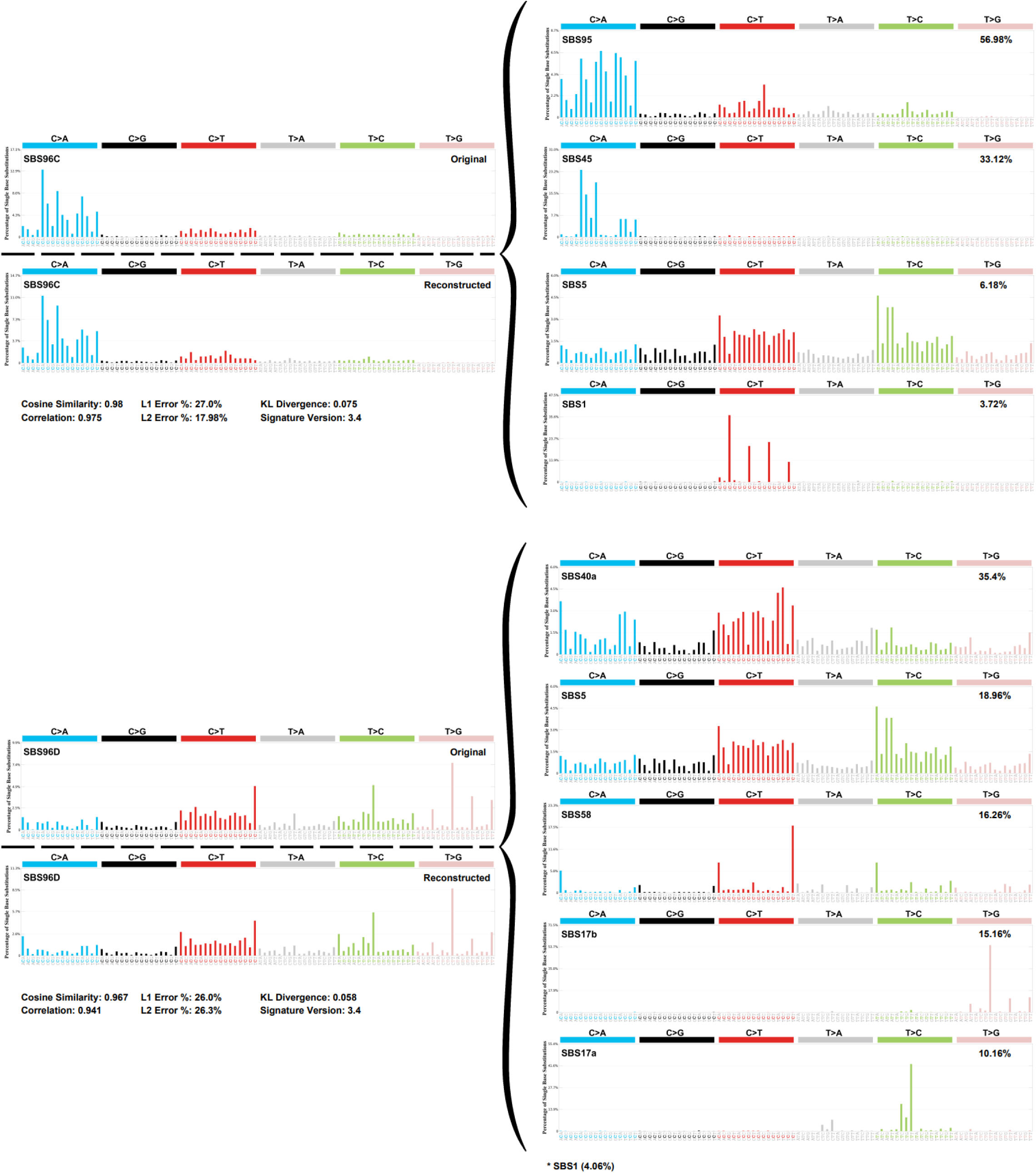
*De novo* SBS mutational signatures extracted from the rat tumors. (**A**) Selection plot of the SigProfilerExtractor tool proposed an extraction solution involving 4 stable signatures. (**B**) Histograms showing the original four *de novo* SBS mutational signatures and their reconstructed versions (left panels). The reconstructed versions were refitted from linear combinations of the known reference signatures (COSMIC, right panels) decomposed from the *de novo* patterns.

**Supplementary Figure S7.**
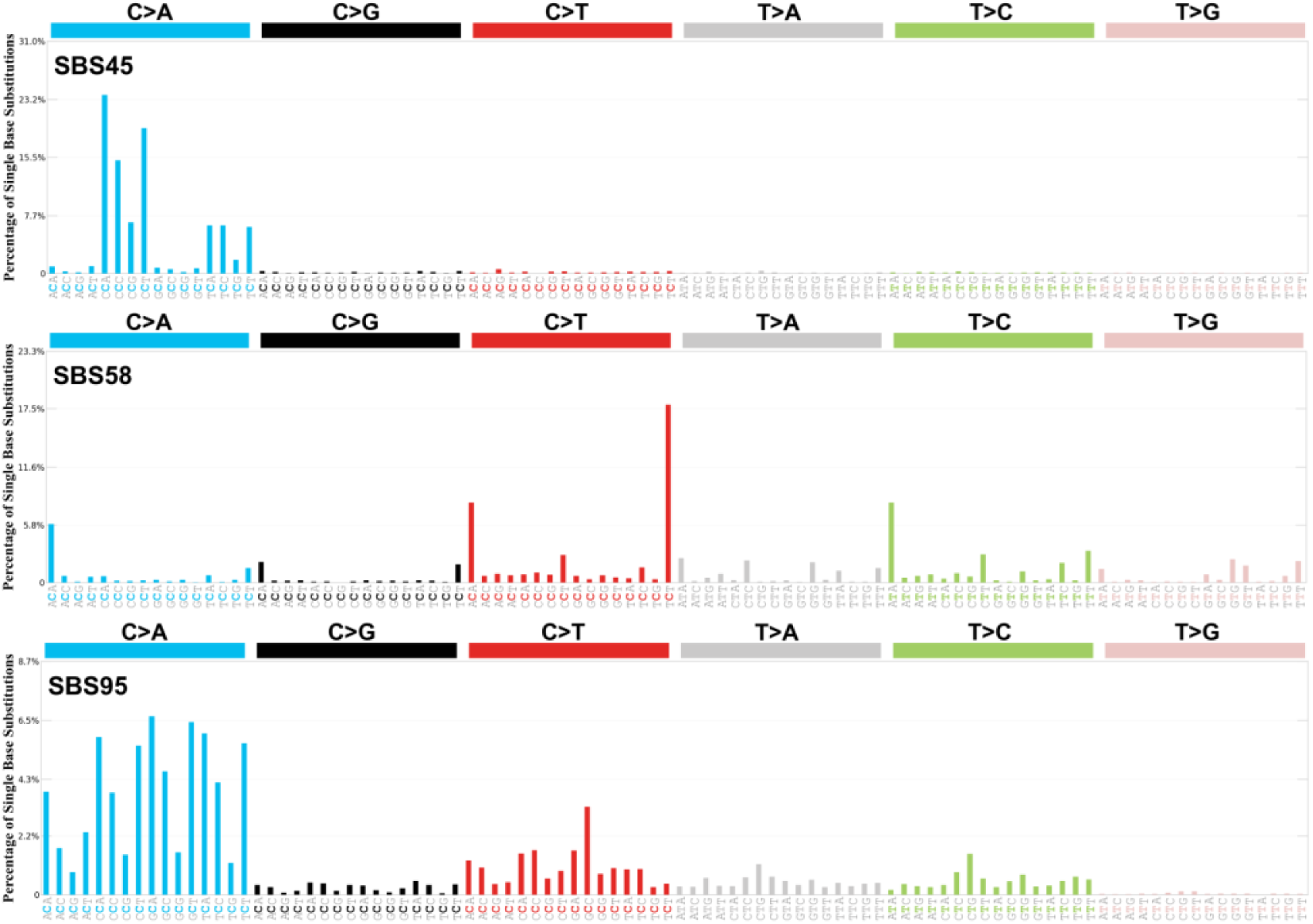
Possible technical-artifact signatures extracted from the rat tumors. COSMIC mutational signatures SBS45, SBS58, SBS95 were occasionally found in some of the rat tumors (see also Fig. 2), possibly representing technical or sequencing artifacts.

**Supplementary Figure S8.**
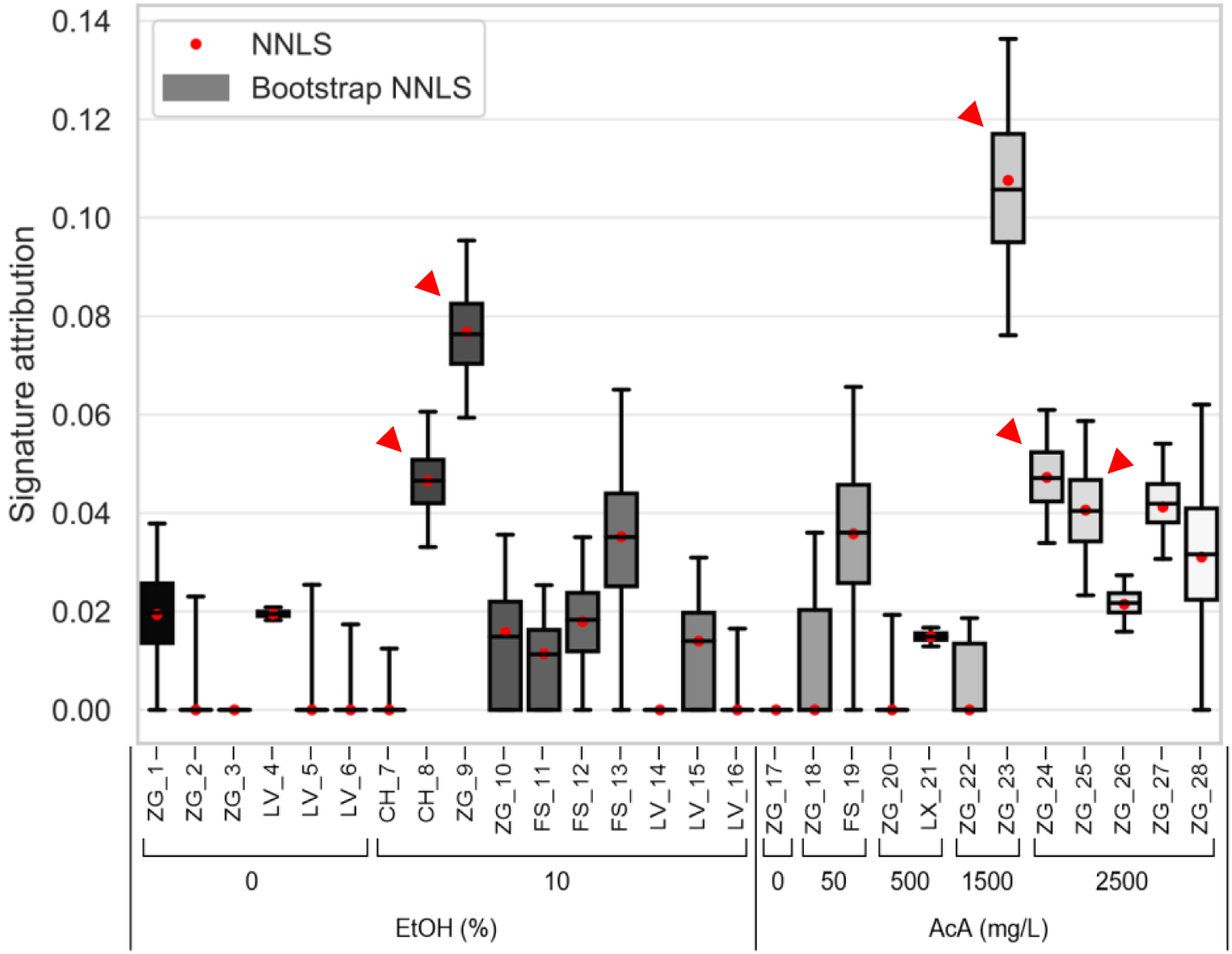
Mutational signature attribution (MSA) of signature SBS17a. Relative attribution of the SBS17a signature – also identified by SigProfilerExtractor - across tumor samples using optimized attribution by the MSA tool. Red arrows – samples for which both SBS17a and 17b were detected by SigProfilerExtractor and MSA (see Fig 2).

**Supplementary Figure S9.**
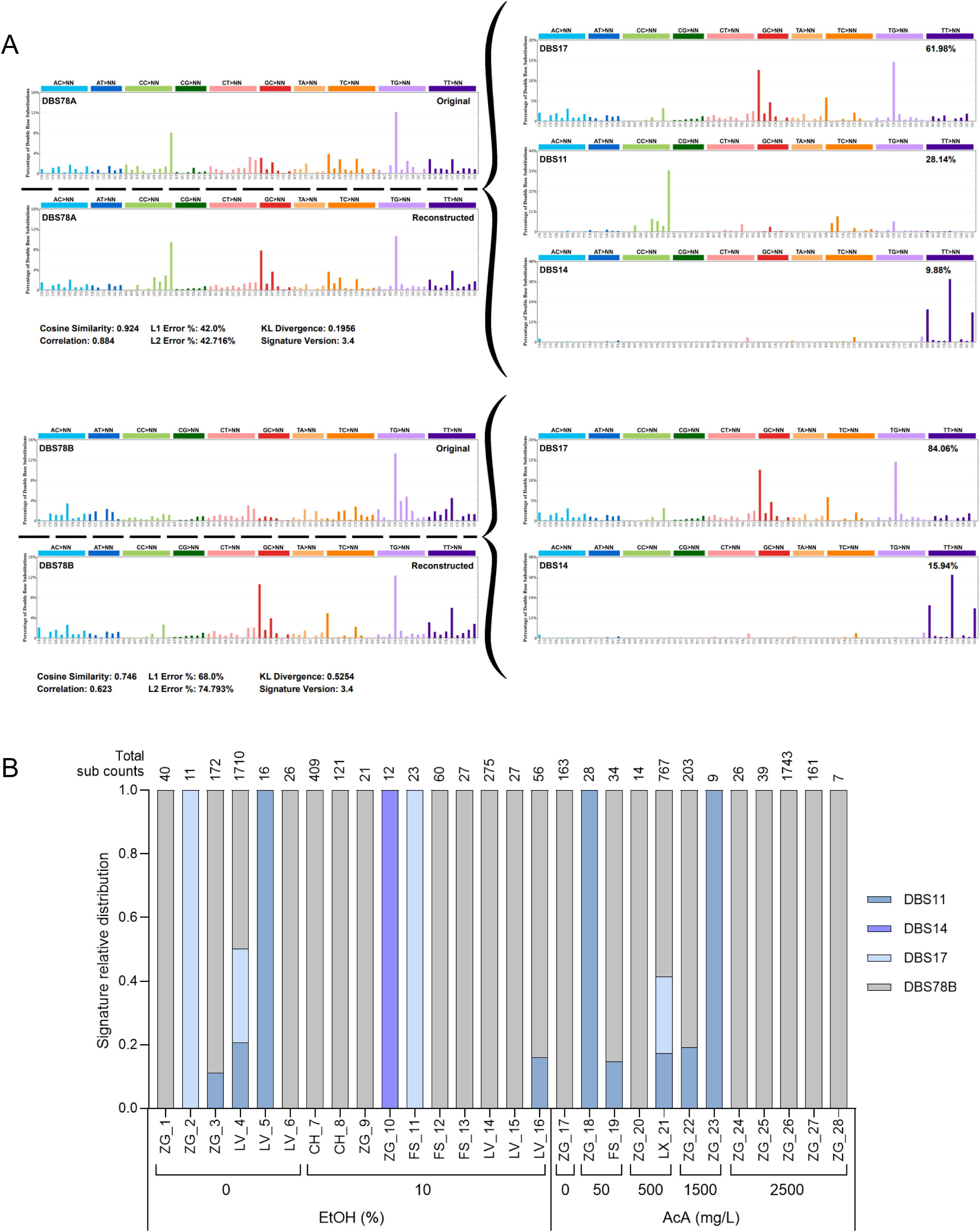
Double base substitution (DBS) mutational signatures in rat tumors. (**A**) Histograms showing the original two *de novo* DBS mutational signatures and their reconstructed versions (left panels). The reconstructed versions were refitted from linear combinations of the known reference signatures (COSMIC, right panels) decomposed from the *de novo* patterns. (**B**) Relative per-sample distribution of the decomposed DBS signatures, including reference COSMIC signatures as well as a *de novo* signature DBS78B not properly decomposed into COSMIC reference signatures. Total DBS mutation counts per sample are shown on top of the graph.

**Supplementary Figure S10.**
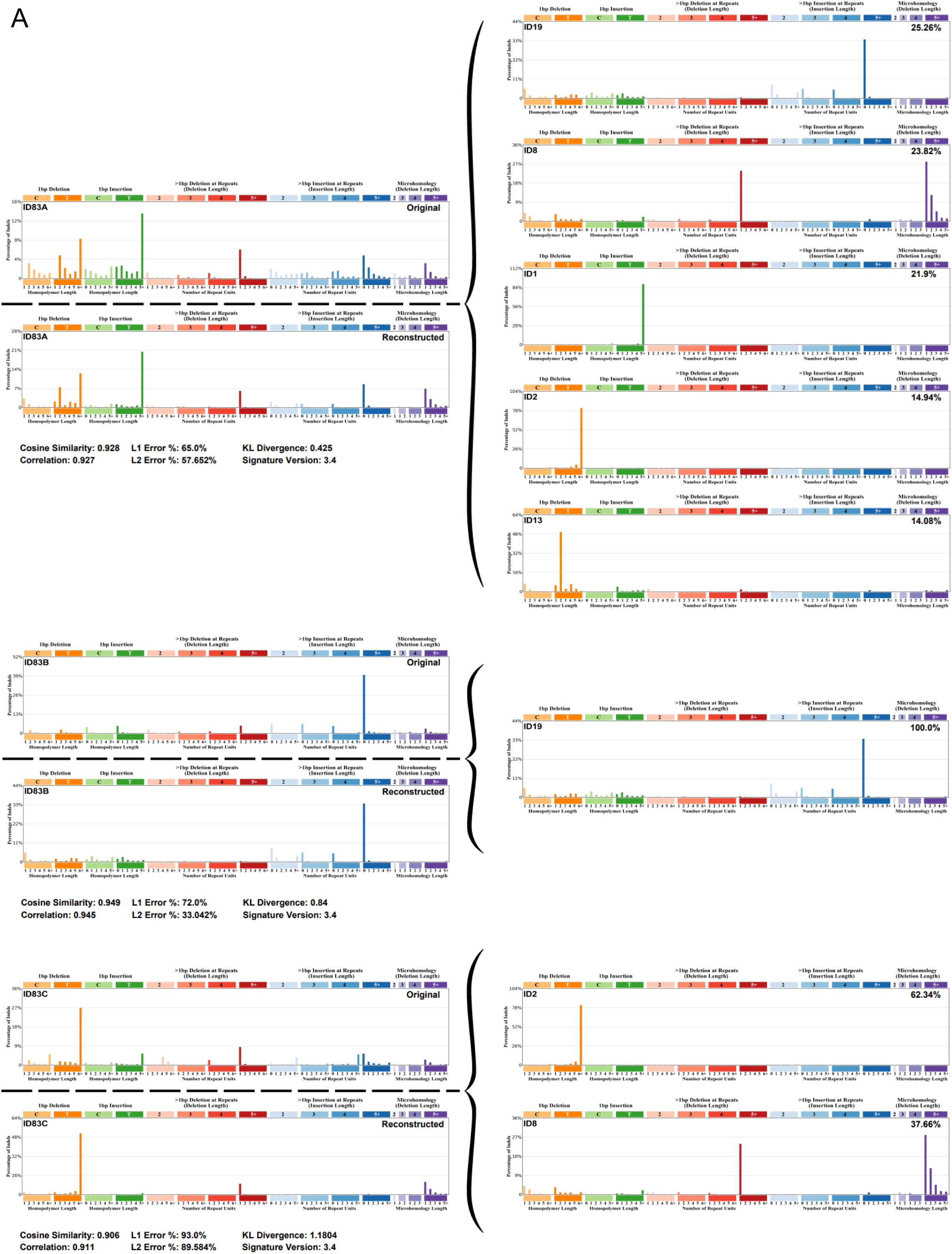
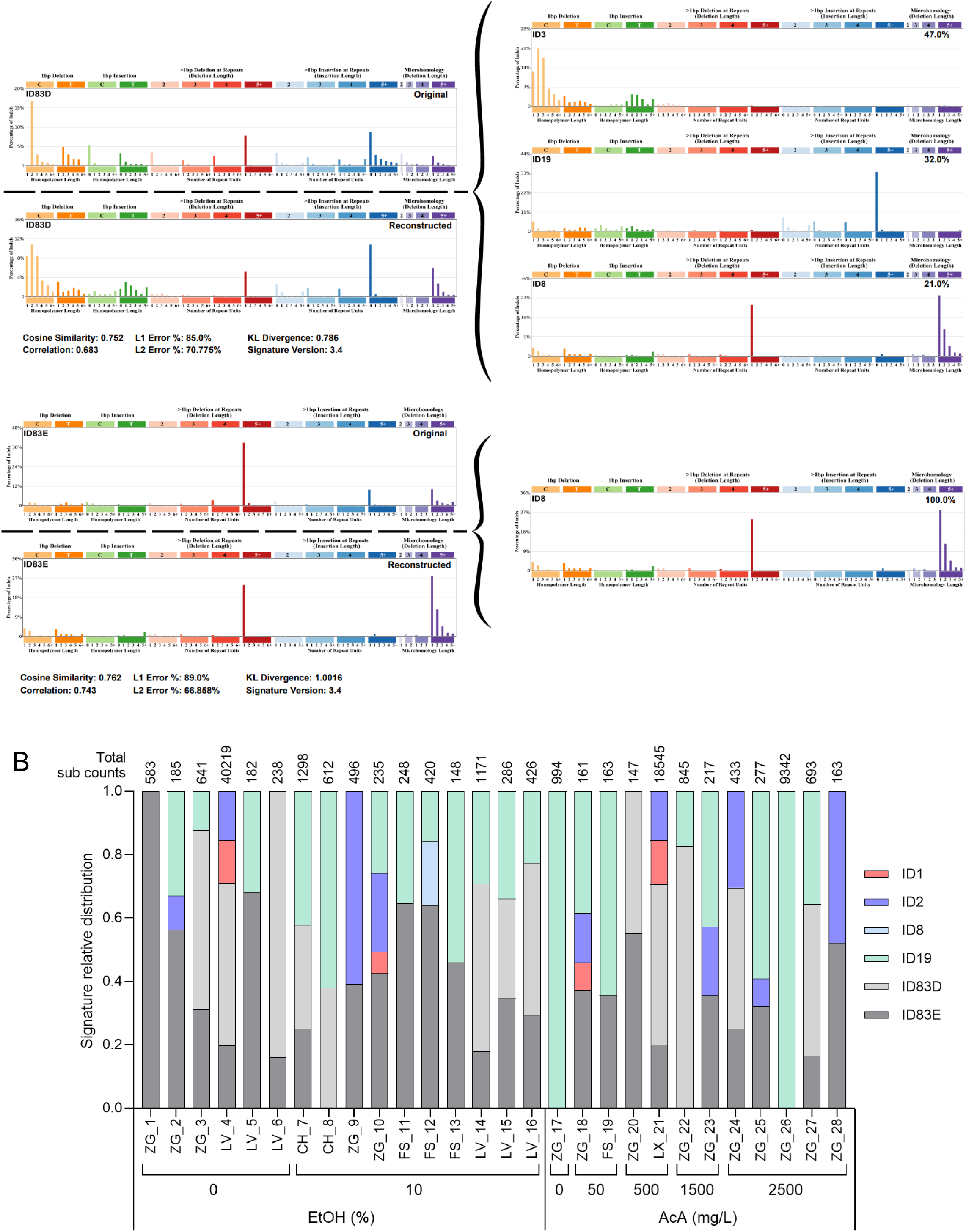
Indel (ID) mutational signatures in rat tumors. (**A**) Histograms showing the original five *de novo* ID mutational signatures and their reconstructed versions (left panels). The reconstructed versions were refitted from linear combinations of the known reference signatures (COSMIC, right panels) decomposed from the *de novo* patterns. (**B**) Relative distribution of the decomposed ID signatures, including reference COSMIC signatures as well as *de novo* signatures ID83D and ID83E not properly decomposed into COSMIC reference signatures. Total ID mutation counts are shown on top of the graph.

**Supplementary Figure S11.**
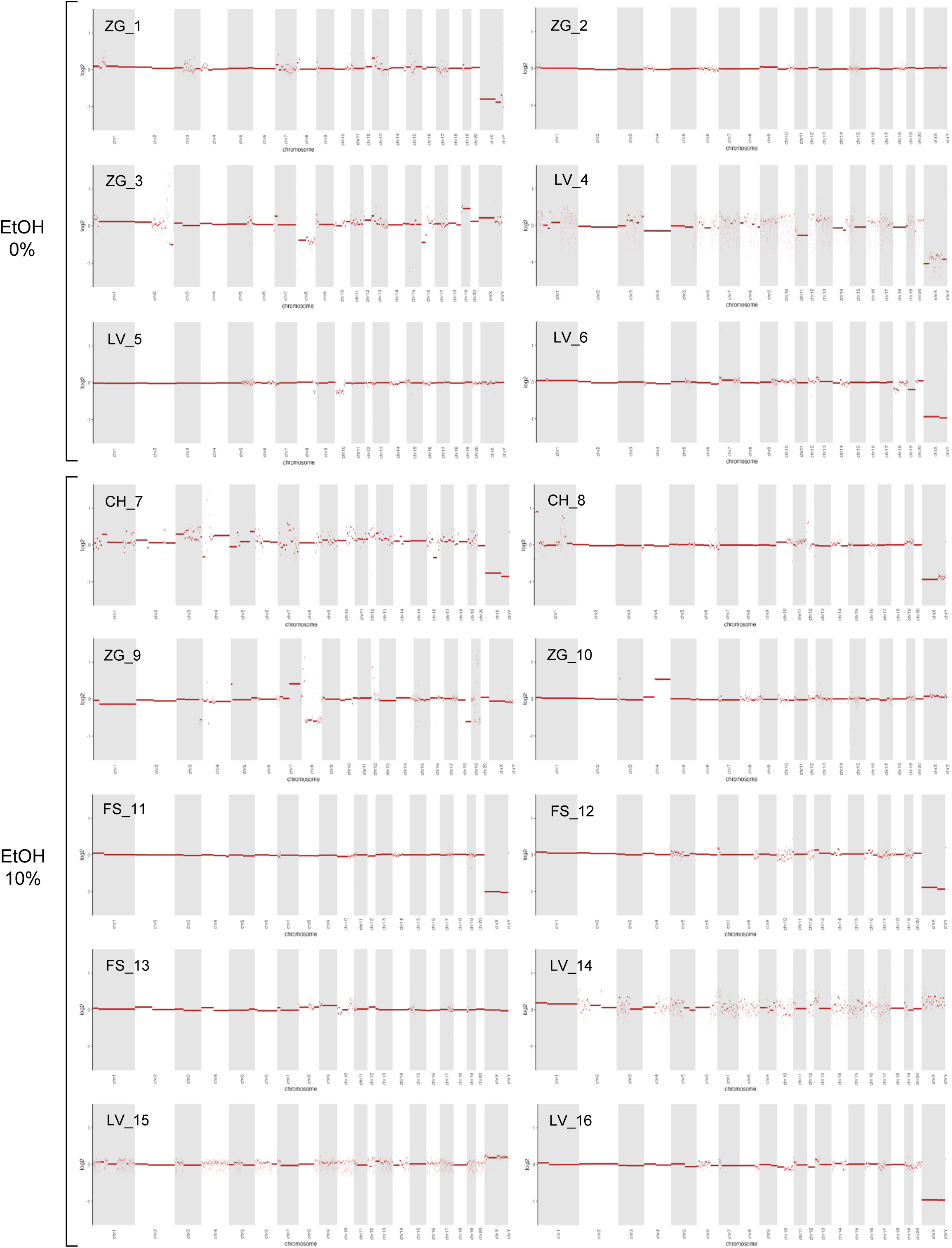
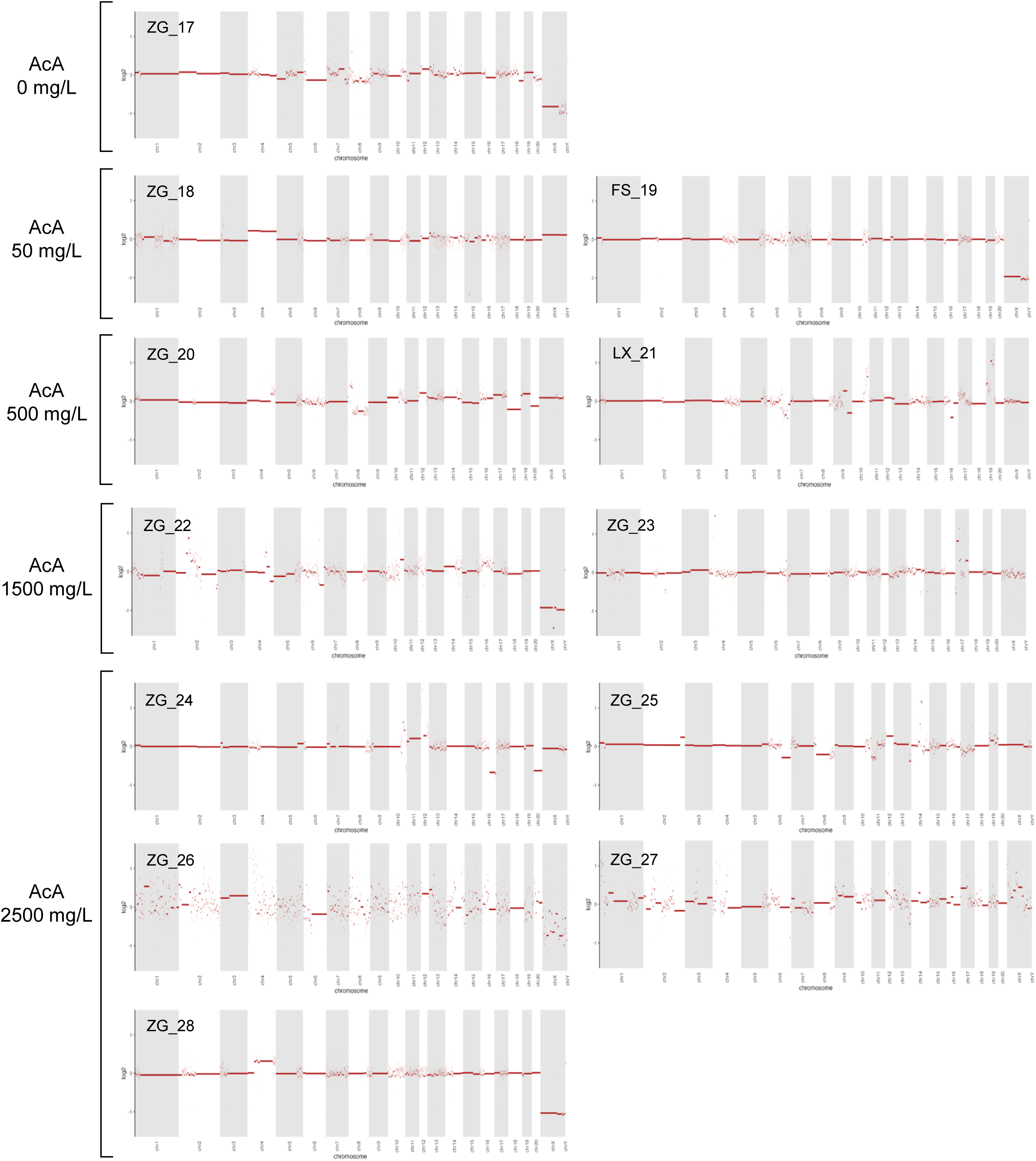
Analysis of copy number alterations (CNA) in the rat tumor samples. Per-sample genome-wide somatic CNA patterns detected in the DNA segments of tumor samples collected on rats exposed to EtOH or AcA at various doses, as indicated. Refer to Methods for additional details.

